# Cbp1, a rapidly evolving fungal virulence factor, forms an effector complex that drives macrophage lysis

**DOI:** 10.1101/2021.09.16.459956

**Authors:** D. Azimova, N. Herrera, L. Duvenage, M. Voorhies, B.C. English, J.C. Hoving, O. Rosenberg, A. Sil

## Abstract

Intracellular pathogens secrete effectors to manipulate their host cells. *Histoplasma capsulatum* (*Hc*) is a fungal intracellular pathogen of humans that grows in a yeast form in the host. *Hc* yeasts are phagocytosed by macrophages, where fungal intracellular replication precedes macrophage lysis. The most abundant virulence factor secreted by *Hc* yeast cells is Calcium Binding Protein 1 (Cbp1), which is absolutely required for macrophage lysis. Here we take an evolutionary, structural, and cell biological approach to understand Cbp1 function. We find that Cbp1 is present only in the genomes of closely related dimorphic fungal species of the Ajellomycetaceae family that lead primarily intracellular lifestyles in their mammalian hosts (*Histoplasma, Paracoccidioides*, and *Emergomyces*), but not conserved in the extracellular fungal pathogen *Blastomyces dermatitidis*. We determine the *de novo* structures of *H*c H88 Cbp1 and the *Paracoccidioides americana* (Pb03) Cbp1, revealing a novel “binocular” fold consisting of a helical dimer arrangement wherein two helices from each monomer contribute to a four-helix bundle. In contrast to Pb03 Cbp1, we show that *Emergomyces* Cbp1 orthologs are unable to stimulate macrophage lysis when expressed in the *Hc cbp1* mutant. Consistent with this result, we find that wild-type *Emergomyces africanus* yeast are able to grow within primary macrophages but are incapable of lysing them. Finally, we use subcellular fractionation of infected macrophages and indirect immunofluorescence to show that Cbp1 localizes to the macrophage cytosol during *Hc* infection, making this the first instance of a phagosomal human fungal pathogen directing an effector into the cytosol of the host cell. We additionally show that Cbp1 forms a complex with Yps-3, another known *Hc* virulence factor that accesses the cytosol. Taken together, these data imply that Cbp1 is a rapidly evolving fungal virulence factor that localizes to the cytosol to trigger host cell lysis.

**Author Summary:** The members of the Ajellomycetaceae fungal family are human pathogens that are responsible for a rising number of mycoses around the world. Calcium binding protein 1 (Cbp1) is a rapidly evolving virulence factor that is present in the genomes of the Ajellomycetaceae species that lead primarily intracellular lifestyles, including *Histoplasma, Paracoccidioides*, and *Emergomyces* but not *Blastomyces*, which remains largely extracellular during infection. Both *Paracoccidioides* and *Histoplasma* Cbp1 homologs are able to cause lysis of macrophages whereas *Emergomyces* homologs cannot. This result is consistent with *Emergomyces africanus* natural infection of macrophages, during which the yeast cells can replicate but cannot actively lyse the host cell. Despite divergence of the primary sequence of *Histoplasma* and *Paracoccidioides* Cbp1 homologs, their protein structures are remarkably similar and reveal a novel fold. During infection, Cbp1 enters the cytosol of the host macrophage, making it the first known virulence factor from an intracellular human fungal pathogen that localizes to the cytosol of the host cell. We also show that Cbp1 forms a complex with another cytosolic virulence factor, Yps-3. Taken together, these studies significantly advance our understanding of *Histoplasma* virulence.

## Introduction

Intracellular pathogens contend with the need to co-opt their host cells to fashion a replicative niche and the requirement to manipulate host-cell survival, sometimes destroying their host cell at the appropriate time to promote exit and further dissemination. To shape their host environment, intracellular pathogens secrete a number of diverse molecules to help the pathogen evade detection by the immune system, scavenge nutrients, and assist in replication and dissemination throughout their host. Bacteria, viruses, parasites, and fungi all secrete effectors, many of which are proteins that are capable of manipulating the biology of the host cell (1-5). Pathogens that reside in nutritionally limiting phagosomal compartments that are normally slated for degradation must be particularly adept at remodeling this niche to make it compatible with microbial replication (6-8). In the case of human pathogenic fungi, we understand much less about secreted effectors of virulence (9-11) compared to bacterial or parasite pathogens. Here we explore the biology of a key virulence effector from the intracellular human fungal pathogen *Histoplasma capsulatum (Hc*).

*Hc* is the causative agent of histoplasmosis and is responsible for high morbidity and mortality even in immunocompetent humans (11-14). This fungus is found world-wide, and in the Central and Eastern United States, *Histoplasma* species are commonly found in the soil of the Ohio and Mississippi River Valleys (15-17). *Hc* is a thermally dimorphic fungus that grows in the soil in a multicellular filamentous form that produces spores known as conidia (18-20). These filamentous fragments and conidia can be inhaled by a mammalian host, where host temperature induces a transition to a pathogenic yeast form. *Hc* yeast are phagocytosed by resident alveolar macrophages in the lungs, where they replicate within a modified phagosomal compartment (21-26). Once *Hc* replicates to high levels, the host cells lyse, thereby releasing yeast cells and facilitating further spread of the fungus to neighboring macrophages.

One of the most abundant yeast-phase proteins secreted by *Hc* into culture supernatants is a small 7.8 kDa protein known as Calcium binding protein 1 (Cbp1) (27-30). As determined previously by an NMR structure and other biochemical analyses, Cbp1 is a dimer with three intramolecular disulfide bridges that make it highly stable (31, 32). The mature secreted form of Cbp1 is 78 amino acids in length, has no known domains, and, prior to this work, had only one known homolog in the closely related fungus *Paracoccidioides*. Interestingly, Cbp1 is a critical virulence factor that is required for virulence of *Hc* in both macrophage and mouse models of infection (29, 33, 34). *Hc* that lacks Cbp1 is able to grow to high levels within the macrophage but is unable to lyse either primary murine bone marrow derived macrophages (BMDMs) or primary alveolar macrophages (33, 34). These data suggest that Cbp1 is a protein effector used by *Hc* to induce lysis of macrophages. Recent work from our lab has demonstrated that Cbp1 actively triggers apoptosis of macrophages through the integrated stress response (ISR), a cascade that is triggered in the cytosol of infected macrophages. The secretion of a functional Cbp1 is absolutely required for this stress response to be initiated in infected macrophages (34). Additionally, we have previously shown that the *Paracoccidioides americana* (Pb03) homolog of Cbp1 is capable of fully restoring lytic capability when expressed in a *Hc cbp1* mutant background (34), suggesting that *Hc* is not the only species that could be using Cbp1 to elicit host cell death. How Cbp1 triggers the ISR and macrophage death is still not fully understood.

Another known secreted effector of *Hc* is the Yeast phase specific 3 (Yps-3) protein (35-37). Like Cbp1, Yps-3 is a secreted factor that is only produced by the pathogenic yeast phase of the fungus and has been previously shown to be important for virulence in a murine model of histoplasmosis using an RNA interference strain with reduced expression of *YPS3* (35). Yps-3 is thought to coat the surface of the yeast cell by binding to chitin (36). Neither Cbp1 nor Yps-3 has been shown to interact with any other fungal proteins.

In this study, we used evolutionary, structural, and cellular approaches to study *Hc* Cbp1 and its homologs. We found new, previously unannotated Cbp1 homologs in the genomes of *Paracoccidioides* and *Emergomyces* species. *Emergomyces* are newly emerging human fungal pathogens that are responsible for a rising incidence of mycoses in immunocompromised humans worldwide (38-41). In contrast, despite its close evolutionary relationship to *Hc*, the extracellular pathogen *Blastomyces* does not harbor a Cbp1 homolog. We determined a new structure of both *Hc* and *Pb* homologs of Cbp1, thereby revealing a novel protein fold. To perform a functional analysis of Cbp1 homologs, we expressed Cbp1 from Pb03 and *Emergomyces* species in the *Hc cbp1* mutant and assessed the ability of each homolog to complement the host lysis defect. Despite the conservation of Cbp1 homologs in the genomes of *Emergomyces* species, these proteins were unlike the Pb03 Cbp1 in that they could not restore lysis during *Hc* infection. For the first time, we assessed the interaction between BMDMs and wild-type *Es. africanus* and discovered that this fungus is capable of robust intracellular replication within BMDMs without any evidence of host-cell lysis, consistent with the inability of *Emergomyces* Cbp1 to complement the *Hc* mutant. Finally, to determine the site of action of *Hc* Cbp1, we discovered that it enters the macrophage cytosol from the *Hc*-containing phagosome, making it the first example of an effector from an intracellular fungal pathogen of humans that accesses the host cell cytosol. Biochemical analyses revealed that Cbp1 forms a complex with Yps-3 and, coupled with our observation that *yps3*Δ mutants cannot achieve maximal lysis of macrophages, suggested the formation of a cytosolic effector complex that mediates host-cell death. Given that Cbp1 1) is unique to the Ajellomycetaceae, 2) shows significant sequence divergence between homologs, and 3) mediates differential lysis between intracellular pathogens, we conclude that the virulence factor Cbp1 is a rapidly evolving protein that is critical for macrophage manipulation by *Hc*.

## Results

### Cbp1 is conserved amongst the Ajellomycetaceae with the exception of *Blastomyces* species

Cbp1 homologs are not broadly present in the fungal kingdom (34). Due to its critical role during intracellular infection, we hypothesized that Cbp1-dependent virulence strategies could be conserved amongst closely related thermally dimorphic fungal species in the Ajellomycetaceae family that are human intracellular pathogens. To identify all putative Cbp1 homologs, we expanded the Cbp1 alignment previously published in (34) by building a Hidden Markov Model (HMM) to search through the annotated protein sets generated from all the additional published genomes of *Emergomyces, Blastomyces*, and *Paracoccidioides* species deposited in GenBank. We detected five Cbp1 *Emergomyces* and two *Paracoccidioides* homologs. To identify the *Emergomyces* homologs, we first defined a syntenic region based on the previously established *Hc* and *Pb* homologs. When we interrogated the corresponding syntenic region *in Emergomyces* species, we uncovered three new *Emergomyces* homologs. All three share the conserved two intron structure found in the *Hc* and *Pb CBP1* genes (28, 34). In the case of *Es. pasteuriana*, we discovered two Cbp1 homologs (*Es. pasteuriana_1* and *Es. pasteuriana_2)*. One homolog was in the syntenic region and maintained the two-intron structure whereas the other was located in a different genomic region and had a three-intron gene structure. Using the *Es. pasteuriana* homologs as templates, we searched through the unannotated *Es. orientalis* genome and found two additional homologs (*Es. orientalis_1, Es. orientalis_2*). As with *Es. pasteuriana*, one homolog was syntenic with a two-intron structure and one was orthologous to the three-intron *Es. pasteuriana_2* Cbp1. In contrast, we found that Cbp1 was not present in any of the *Blastomyces* genomes. Unlike the other three major members of this fungal family, *Blastomyces* leads a largely extracellular lifestyle *in vivo (41-43)*, suggesting that Cbp1 could be a virulence factor that represents a specialized adaptation to an intracellular lifestyle inside of a mammalian host.

We aligned the new Cbp1 homologs to the Cbp1 consensus sequence generated by the HMM (Fig. 1A) along with the previously characterized *Hc* and *Pb03* Cbp1 homologs, which have been previously shown to be necessary for macrophage lysis (29, 33, 34). The 5’ portion of the Cbp1 coding sequence, which contains the putative signal peptide necessary for extracellular secretion of the protein, is well conserved amongst all the homologs. However, the residues flanking the putative cleavage site of the signal sequence, as based upon *Hc* G217B Cbp1, are poorly conserved. The N-terminus of the mature protein is fairly well conserved whereas the latter half of the predicted coding sequence is highly conserved, especially the six cysteines that form the three intramolecular disulfide bridges (31, 32).

**Fig. 1.**
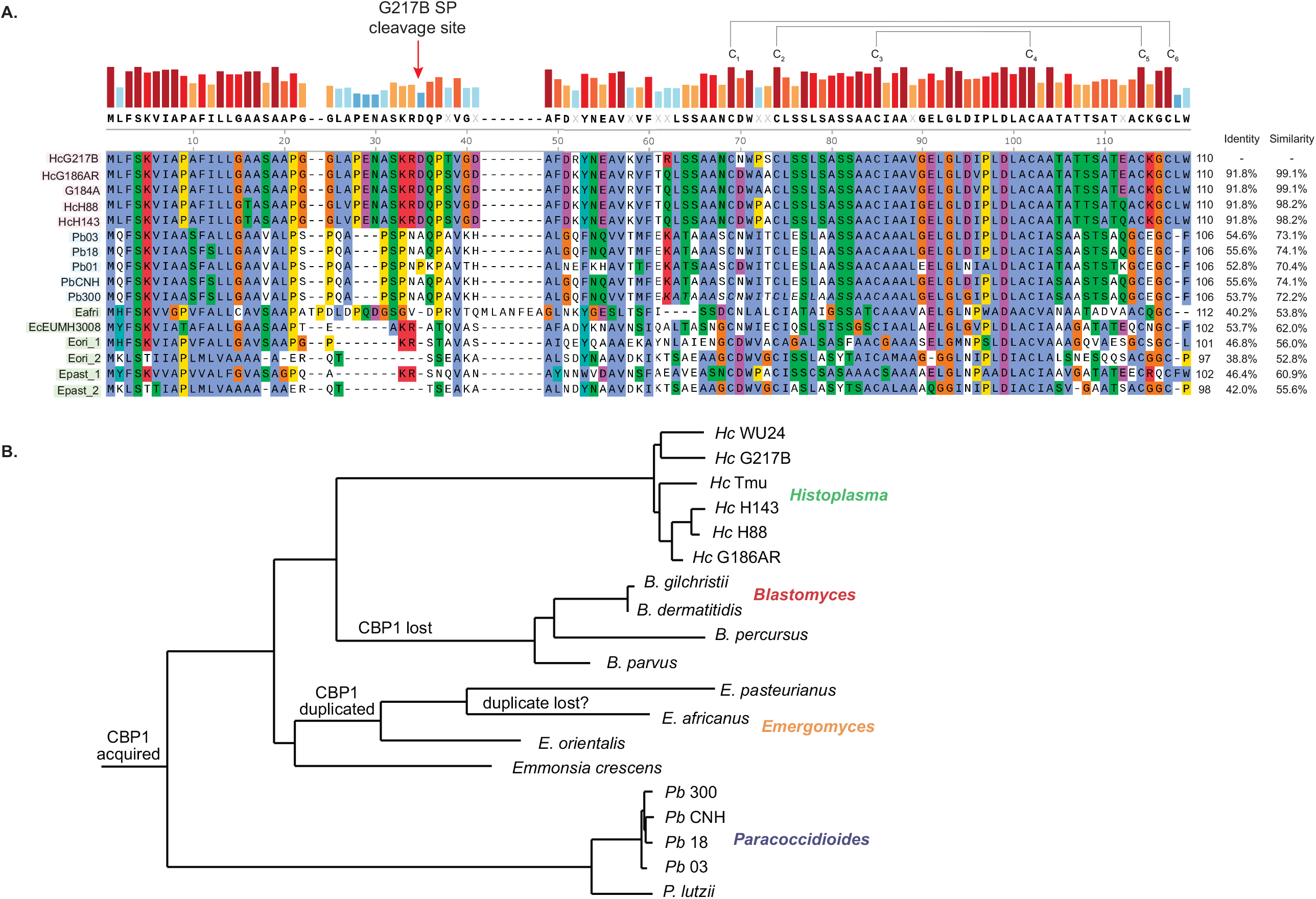
Cbp1 is conserved among closely related thermally dimorphic fungi that are intracellular pathogens of humans. A. Protein alignments of *Hc* G217B Cbp1 and closely related homologs. The site of signal peptide cleavage in the *Hc* G217B protein as well as the six cysteine residues that form intramolecular disulfide bonds are highlighted. The sequences of three *Hc* homologs (G217B, H88, G186AR), one *Pb* homolog (Pb03), one *Es. africanus* homolog, one *E*.*crescens* homolog, two *Es. orientalis* homologs, and two *Es. pasteurianus* homologs are shown. The percentages of identity and similarity of each homolog relative to the *Hc* G217B Cbp1 are listed. B. Species tree that tracks the acquisition and loss events of Cbp1 in Ajellomycetaceae species that are human fungal pathogens.

When comparing the sequences of the Cbp1 homologs on a primary amino acid level relative to the sequence of G217B Cbp1, it is clear that these sequences diverge rapidly. We see that the Cbp1 sequences within each genus (*Histoplasma, Paracoccidioides*, and *Emergomyces*) are much closer to each other than they are to homologs from a different genus. The percent identity between G217B and G186AR/H88 Cbp1 is high, at 91.8% (Fig. 1A). The next closest sequence to G217B is that of Pb03 and other *Paracoccidioides* species, which range between 52-55% identical (Fig. 1A). *Emergomyces* Cbp1 homologs show less identity to G217B Cbp1, ranging between 38-53% (Fig. 1A). Using Phylogenic Analysis by Maximum Likelihood (PAML) (44) over the alignment of the 11 orthologs found in the syntenic region of the G217B *Hc* Cbp1, we find that there is significant support for a positive selection model (M8/M7) of Cbp1 in these genomes with a p = 7.918e-06 (44).

The most parsimonious explanation for the emergence of Cbp1 in this fungal family is that Cbp1 evolved in an early common ancestor and was then subsequently lost from the *Blastomyces* ancestral species as it diverged away from the other members of Ajellomycetaceae (Fig. 1B). Similarly, the duplication of the Cbp1 gene in *Es. pasteuriana* and *Es. orientalis* may have occurred in a common ancestor of *the Es. orientalis, Es. pasteuriana*, and *Es. africanus* species. We note that this implies either subsequent loss of the Cbp1 paralog in *Es. africanus* or omission of this gene from the *Es. africanus* assembly (GCA_001660665.1), which is highly fragmented and 3MB smaller than the other *Emergomyces* genomes. Taken together, these data suggest that Cbp1 is a rapidly evolving protein that was gained recently amongst the fungi of the Ajellomycetaceae, diverged rapidly between the various species in this family, and remains under positive selection.

### Structures of Hc H88 and Pb03 Cbp1 reveal a novel helical “binocular” fold

To determine if there is any structural similarity between these homologs and the previously published G186AR NMR Cbp1 structure (31), we used X-ray crystallography to solve the structure of a subset of homologs. Since Cbp1 makes up the vast majority of the *Hc* secretome (45), we purified the protein directly from culture supernatants with minimal perturbation of the native protein. We used a similar approach to purify *Hc* G217B, *Hc* G186AR, *Hc* H88, Pb03, *E. crescens, and Es. africanus* Cbp1 homologs expressed in *Hc* (Fig. S1). Crystals of *Hc* H88 and Pb03 Cbp1 homologs formed after a week and diffracted to 1.5 Å and 1.6 Å respectively (Fig. 2A). To resolve the phases, a heavy atom derivative was prepared by soaking the Pb03 crystals in K_2_PtCl_4_. Two datasets were collected at the L III Pt-edge at 11562 eV and at a remote energy of 13500 eV. Multiple anomalous diffraction (MAD) data were used to search for the heavy atom positions using the Phaser program in the Phenix crystallography programming suite. The structure of Pb03 was truncated at C-terminal residues 65-67 from chain A and 67-68 from chain B, as the remaining amino acids were not resolved in the refined density. The final refinement statistics were Rwork/free= 0.21 / 0.22 and the data were deposited under PDBID 7R6U (Table 1). The diffraction data from *Hc* H88 was resolved the by molecular replacement using the Pb03 model. Using this model in the Phaser program in the Phenix crystallography programming suite, we obtained a TFZ score of 8.4 and an LLG score of 63.271, supporting the validity of the molecular replacement solution. The final refinement statistics were Rwork/free = 0.21 / 0.23 and the data were deposited under PDBID 7R79 (Table1).

**Fig. 2.**
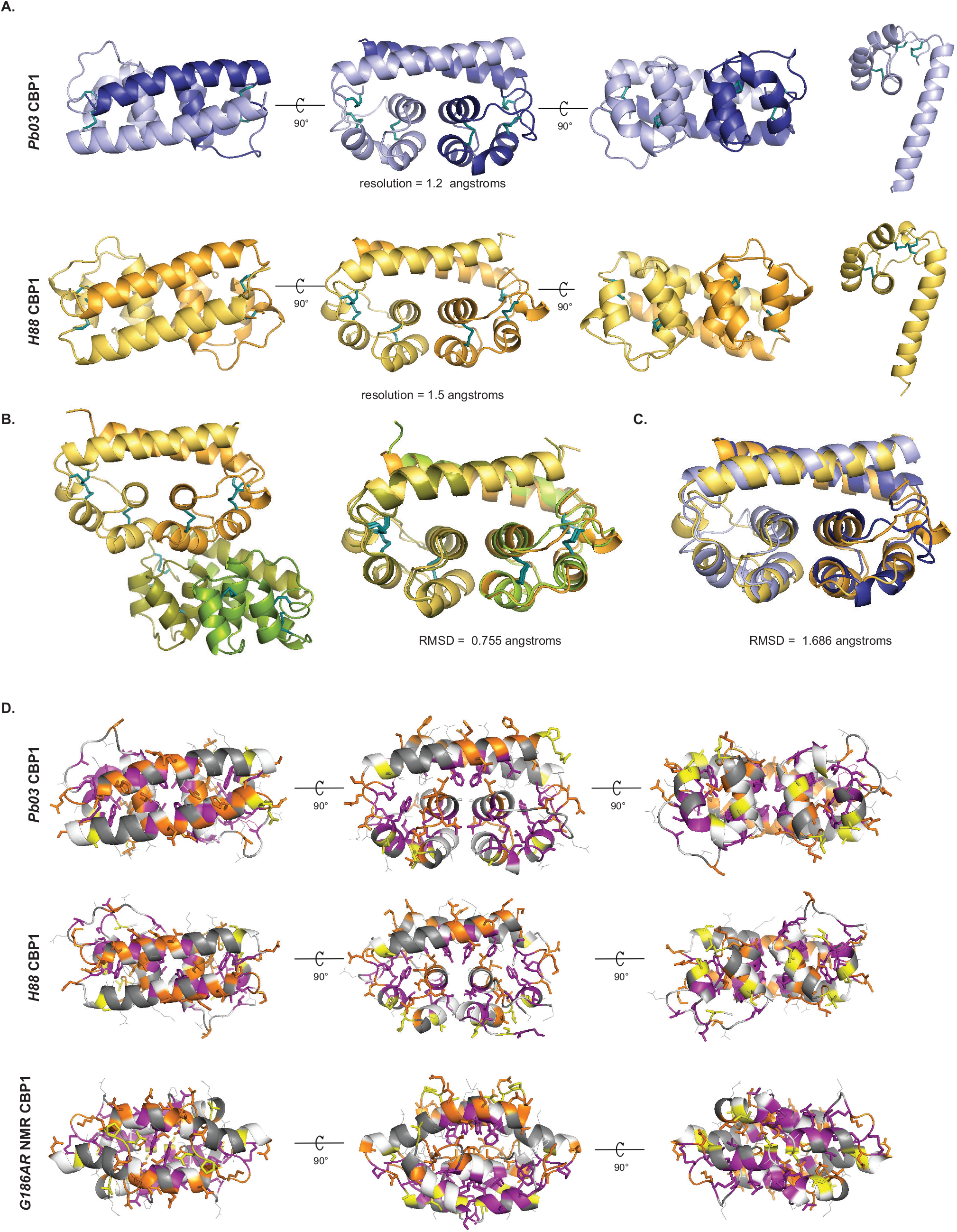
Crystal structures of H88 and Pb03 Cbp1 reveal a novel “binocular” fold. A. Structures of *Pb03* and *H88* Cbp1 as determined by crystallographic methods are shown in native dimer form and as a monomer. B. Quaternary structure of the H88 Cbp1 shows a dimer of dimers that interact though the C-terminal helical bundles. The two dimers in the H88 quaternary structure are aligned and the RMSD score was determined to be 0.755 angstroms. C. Structural alignments of *Pb03* and *H88* Cbp1 structures reveal a highly similar fold with an RMSD value of 1.686 angstroms. D. Prior experiments in our laboratory determined that the purple residues are required for secretion, the orange residues are absolutely required for lysis, and the yellow residues contribute to maximal lysis (34). When these residues are modeled on the *Pb03* or *H88* crystal structure (top two rows), the purple residues face inwards and the orange and yellow residues are predicted to be on the surface of the protein facing outwards. In contrast, modeling the purple, orange, and yellow residues on the NMR structure does not result in a consistent pattern.

**Table 1.** Data collection and refinement statistics.

|  | **H88 Cbp1** | **Pb03 Cbp1** | **Pb03 Cbp1- E1** | **Pb03 Cbp1- E2** |
| --- | --- | --- | --- | --- |
| **Wavelength** | 1.11 | 1.11 | 1.07 | 0.90 |
| **Resolution range** | 42.27 - 1.6<br>(1.657 - 1.6) | 44.14 - 1.55<br>(1.605 - 1.55) | 43.95 - 3.0<br>(3.108 - 3.0) | 43.89 - 3.0<br>(3.107 - 3.0) |
| **Space group** | C 1 2 1 | P 41 21 2 | P 41 21 2 | P 41 21 2 |
| **Unit cell** | 88.55 46.697 71.255<br>90 109.097 90 | 46.507 46.507<br>140.123 90 90 90 | 46.299 46.299<br>139.775 90 90<br>90 | 46.231 46.231<br>139.716 90 90<br>90 |
| **Total reflections** | 240007 (24131) | 559121 (43919) | 323980 (31184) | 324834 (35562) |
| **Unique reflections** | 36276 (3571) | 23244 (2261) | 3415 (334) | 3408 (336) |
| **Multiplicity** | 6.6 (6.8) | 24.1 (19.4) | 94.9 (93.4) | 95.3 (105.8) |
| **Completeness (%)** | 99.32 (98.29) | 99.92 (99.56) | 99.85 (100.00) | 99.85 (100.00) |
| **Mean I/sigma(I)** | 21.74 (1.83) | 18.12 (0.50) | 32.41 (6.35) | 40.10 (8.52) |
| **Wilson B-factor** | 25.49 | 24.85 | 66.00 | 67.64 |
| **R-merge** | 0.04273 (0.9169) | 0.1218 (4.973) | 0.2405 (1.501) | 0.1782 (1.136) |
| **R-meas** | 0.04643 (0.9925) | 0.1244 (5.107) | 0.2418 (1.509) | 0.1792 (1.142) |
| **R-pim** | 0.01792 (0.3761) | 0.0253 (1.142) | 0.0245 (0.1551) | 0.01811 (0.11) |
| **CC1/2** | 1 (0.758) | 1 (0.219) | 1 (0.972) | 1 (0.989) |
| **CC\*** | 1 (0.929) | 1 (0.6) | 1 (0.993) | 1 (0.997) |
| **Anomalous completeness** | - | - | 100.0 (100.0) | 100.0 (100.0) |
| **Anomalous multiplicity** | - | - | 55.5 (51.3) | 55.7 (58.3) |
| **DelAnom correlation** | - | - | 0.934 (0.265) | 0.948 (0.313) |
| **Mid-Slope of Anom Normal** | - | - | 1.889 | 2.147 |
| **Reflections used in refinement** | 36270 (3571) | 23226 (2251) | - | - |
| **Reflections used for R-free** | 1473 (145) | 1129 (106) | - | - |
| **R-work** | 0.2136 (0.3079) | 0.2101 (0.3366) | - | - |
| **R-free** | 0.2352 (0.3074) | 0.2238 (0.3334) | - | - |
| **CC(work)** | 0.951 (0.788) | 0.959 (0.614) | - | - |
| **CC(free)** | 0.944 (0.717) | 0.961 (0.716) | - | - |
| **Number of non-hydrogen atoms** | 2333 | 1145 | - | - |
| **macromolecules** | 2146 | 1048 | - | - |
| **ligands** | 0 | 0 | - | - |
| **solvent** | 187 | 97 | - | - |
| **Protein residues** | 306 | 152 | - | - |
| **RMS(bonds)** | 0.006 | 0.008 | - | - |
| **RMS(angles)** | 0.87 | 0.99 | - | - |
| **Ramachandran favored (%)** | 97.32 | 97.97 | - | - |
| **Ramachandran allowed (%)** | 2.68 | 2.03 | - | - |
| **Ramachandran outliers (%)** | 0.00 | 0.00 | - | - |
| **Rotamer outliers (%)** | 0.00 | 0.00 | - | - |
| **Clash score** | 2.62 | 0.00 | - | - |
| **Average B-factor** | 32.83 | 35.31 | - | - |
| **macromolecules** | 32.80 | 34.62 | - | - |
| **solvent** | 44.19 | 42.71 | - | - |
| **PDB ID** | 7R79 | 7R6U | - | - |

Statistics for the highest-resolution shell are shown in parentheses.

The asymmetric unit for the Pb03 Cbp1 crystal contained a single dimer, whereas the asymmetric unit for the H88 crystal contained a dimer of dimers (Fig. 2A and 2B). An alignment of the two dimers of H88 Cbp1 structure gave a root mean squared deviation (RMSD) value of 0.755 Å, indicating the structure remains largely unchanged amongst the dimers (Fig. 2B). An alignment of the Pb03 and the *Hc* CBP1 dimers gave an RMSD of 1.6 Å, indicating the structures of the distinct homologs are conserved (Fig. 2C). These structures had no significant structural homology to any known proteins according to DALI (46) or VAST (47) searches but are highly similar to each other as can be seen when the dimers of Pb03 and H88 are aligned (Fig. 2C). The Cbp1 structures displayed an alpha-helical dimer arrangement with an intermolecular four-helix bundle fold at the core formed from the 3^rd^ and 4^th^ helices of each monomer, creating a binocular shape, and thus we dubbed this new arrangement as a “binocular” fold. The bundle formed by two of the three C-terminal helices of each monomer contains a number of aliphatic residues (Ile55, Pro56, Leu59, Ala63) that pack against the longer N-terminal helices (Fig. S2A). The N-terminal helices from both monomers interact with each other in an anti-parallel manner and their register relative to each other appears to be determined by the N-terminal residues packing against the aliphatic patch underneath them. For both Pb03 and H88 Cbp1, the three disulfide bonds present in each monomer orient the alpha helices so that the aromatic residues (Phe12, Phe19, Trp30) are packed in the interior of the protein, creating the hydrophobic core (Fig. S2A).

The Pb03 and H88 structures were similar when they were aligned, yielding an RMSD of 1.6 Å (Fig. 2C). An emerging property of the Cbp1 structures is the conservation of negative charge over various surfaces of the protein at neutral pH (Fig. S2C) (48, 49). The Pb03 structure has a clear negative charge at the C-terminal end of the protein and was partially positive at the interface between the N-terminal helices. The H88 structure was not as clearly polarized but did display a groove of strong negative charge on its C-terminal side. These charges were clearly visible in both the dimers and individual monomers, indicating intrinsic properties of the monomer of the protein (Fig. S2C). When the tertiary structures of *Emergomyces* homologs were modelled (50) on the Pb03 Cbp1 backbone, the variation in surface charge was evident (Fig. S2D).

The previously published *Hc* G186AR NMR Cbp1 structure (31) is distinct from the X-ray crystallography structures determined in this work. When the new structures were aligned against the G186AR NMR structure, the resulting RMSD value was 5.6 Å and 5.8 Å for Pb03 and H88 Cbp1, respectively (Fig. S2B), consistent with the NMR structure not being found by DALI and VAST searches of the crystal structures. In this alignment, a major difference lies in the orientation of the three C-terminal alpha-helices in the crystal structure vs. the NMR structure. In the crystal structure, the C-terminal helices were perpendicular to the N-terminal helices, whereas in the NMR structure they are parallel. In the crystal structure, the N-terminal helices were naturally more linear, resulting in an alignment in a different register relative to each other, and suggesting alternate helix-helix interaction pairs. Finally, in the NMR structure, the two monomers are intercalated, whereas in the crystal structures this was not the case.

In prior work, we used alanine scanning to identify Cbp1 residues that are critical for function of G217B Cbp1 (34) (Fig. 2D). The alanine scan identified residues that, when mutated to alanine, resulted in Cbp1 variants which were (1) incapable of being secreted, (2) capable of being secreted but only caused partial macrophage lysis, or (3) capable of being secreted but completely deficient in macrophage lysis capability. Notably, when we modeled these residues on the Pb03 or H88 crystal structure, we found that the residues that were required for Cbp1 protein secretion were all facing inwards away from the protein surface, suggesting they are important for proper folding (Fig. 2D). This observation was consistent with a role in proper packing of the hydrophobic core. The majority of the residues that were necessary or contributed to Cbp1 lytic capability had side chains that were oriented out towards the external surface and were preferentially located in the N-terminal helix, suggesting that those residues create surfaces that are necessary for Cbp1 function. We did not observe concordance between the alanine mutant phenotypes and the location of the corresponding residues on the *Hc* G186AR NMR structure (Fig. 2D). Thus, the alanine scan data is highly consistent with the newly determined crystal structures of *Hc* H88 Cbp1 and its homologs.

### *Emergomyces* Cbp1 homologs cannot complement the *Hc cbp1* mutant

To further probe the function of the newly identified Cbp1 homologs, we expressed them in *Hc* to determine whether they could complement the *cbp1* mutant. The *Hc* (G217B) *cbp1* mutant is unable to lyse macrophages after intracellular infection (33, 34). A second phenotype of the *Hc cbp1* mutant is a delay in intracellular growth after macrophage infection, although ultimately the *cbp1* mutant reaches high levels of intracellular fungal burden equivalent to or exceeding that of wild-type *Hc* (33, 34). We have previously shown that expression of the *Paracoccidioides americana* (Pb03) Cbp1 homolog in the *Hc* (G217B) *cbp1* mutant can restore macrophage lysis (34). We expressed five of the newly detected *Emergomyces* Cbp1 homologs (*Es. africanus, Es. orientalis_1, E. crescens, Es. pasteuriana_1* and *Es. pasteuriana_2 Cbp1*) in the *Hc* G217B *cbp1* background to see if these homologs can complement either the ability of *Hc* to lyse macrophages or grow intracellularly without delay. To monitor for production and secretion of each of the *Emergomyces* homologs, we subjected concentrated culture supernatants from multiple isolates expressing each of the Cbp1 homologs to SDS-PAGE to monitor for the presence of a prominent Cbp1 band. We found that all five homologs were found in *Hc* culture supernatants in comparable amounts when expressed under the control of the *Emergomyces* native signal peptide (Fig. 3A). Their identity was also confirmed with mass spectrometry of chymotrypsin digested peptides from SDS-PAGE gel bands.

**Fig. 3.**
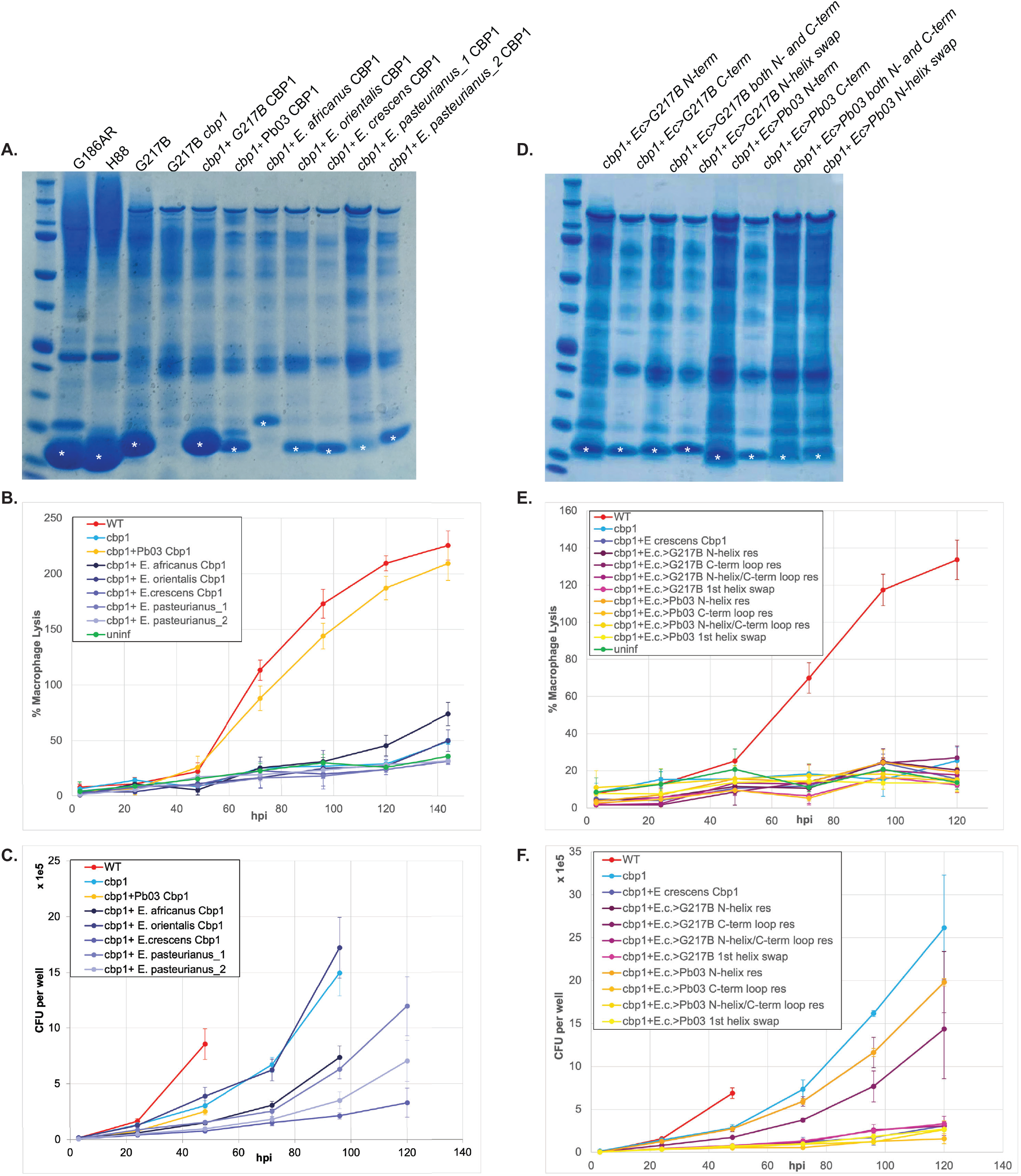
*Emergomyces* species contain Cbp1 homologs in their genomes that can be expressed and secreted by the *Hc* G217B strain but are insufficient to confer macrophagelysis. A. InstantBlue™ stained SDS-PAGE gel of culture supernatants of *Hc* G186AR, *Hc* H88, *Hc* G217B, *Hc* G217B *cbp1* mutant, and *Hc* G217B *cbp1* mutant expressing the G217B, *P. americana (Pb03), E. crescens, Es. orientalis, Es. pasteurianus_1, Es. pasteurianus_2, or Es. africanus* Cbp1. White asterisk denotes the bands that were excised for mass spectrometric analysis to confirm their identity as the appropriate Cbp1. B. *Hc cbp1* mutant isolates expressing the *Emergomyces* Cbp1 homologs were used to infect BMDMs at an MOI of 1 as described in the Methods. “WT” indicates the *Hc* G217B *ura5* mutant carrying the *URA5* gene on a plasmid. LDH release was used to quantify percent host cell lysis. C. Intracellular growth of *Hc cbp1* mutant isolates expressing the *Emergomyces* Cbp1 homologs was determined from BMDM infection as described in the Methods. “WT” indicates the *Hc* G217B *ura5* mutant carrying the *URA5* gene on a plasmid. At 48 hpi, CFU counts were terminated for the WT *Hc* infection because the extent of host-cell lysis made it difficult to distinguish between intracellular and extracellular growth of *Hc*. D. InstantBlue™ stained SDS-PAGE gel of culture supernatants of G217B *Hc cbp1* mutant expressing the chimeric constructs that convert key residues from *E. crescens* Cbp1 into their counterpart residue from G217B or Pb03 Cbp1 in the N-terminus, the C-terminal loops, both the N-terminus and C-terminal loops, or a 1^st^ helix swap. E. *Hc cbp1*-mutant isolates expressing *E. crescens* chimeric Cbp1 homologs were used to infect BMDMs and LDH release was quantified. F. Intracellular growth of *Hc cbp1* mutant isolates expressing the *E. crescens* chimeric Cbp1 homologs was determined from BMDM infection as described in the Methods. “WT” indicates the *Hc* G217B *ura5* mutant carrying the *URA5* gene on a plasmid. At 48 hpi, CFU counts were terminated for the WT *Hc* infection because the extent of host-cell lysis made it difficult to distinguish between intracellular and extracellular growth of *Hc*.

To determine if expression of any of these homologs could complement the phenotypes of the *Hc* G217B *cbp1* mutant, we infected primary murine bone marrow-derived macrophages (BMDMs) with our cohort of heterologous expression strains and monitored for the release of lactate dehydrogenase into the supernatant as a measure of host cell lysis. The *Hc cbp1* mutant fails to lyse BMDMs and also exhibits an intracellular growth delay before it undergoes replication during infection. The Pb03 homolog was able to fully restore the lytic capability of the *cbp1* mutant (Fig. 3B) but did not complement the growth delay as measured by intracellular CFU counts (Fig. 3C). Surprisingly, none of the *Emergomyces* Cbp1 homologs were able to restore the ability of the *Hc* G217B *cbp1* mutant to lyse macrophages (Fig. 3B). The *Emergomyces* homologs showed variable ability to support intracellular growth in macrophages: *cbp1* mutant cells expressing the *Es. orientalis* Cbp1 grew on par with the *Hc cbp1* mutant, cells expressing the *E. crescens* Cbp1 never grew well inside macrophages, and the *Es. africanus, Es. pasteurianus-1 and -2* alleles conferred variable growth (Fig. 3C), suggesting some of these variants could interfere with intracellular growth.

To assess which molecular differences between Cbp1 homologs might correlate with differences in function, we created a series of chimeric Cbp1 protein constructs (Fig. S3A-B). Of the *Emergomyces* Cbp1 homologs, we chose *E. crescens* Cbp1 as a template because it had the highest percent identity to *Hc* G217B Cbp1. Based on the alanine scan data described above, we identified key amino acids in *Hc* G217B Cbp1 that are required for macrophage lysis (34) (Fig. S3A). A subset of the analogous amino acids in *E. crescens* Cbp1 were changed to the corresponding G217B *Hc or* Pb03 Cbp1 residues. These residues were selected based on the following criteria: whether their sidechains were facing the external surface, whether they were necessary for lysis based on the alanine scan data, and whether there was a significant change in either polarity, charge, or size as compared to *Hc* G217B Cbp1. Four chimeric constructs were ultimately generated that either exchanged the 1) entire N-terminal helix, 2) only the residues necessary for lysis in the N-terminal helix, 3) the residues necessary for lysis in the loops and helices of the C-terminal helical bundle, or 4) a combination of constructs 2 and 3 (Fig. S3A). These chimeric Cbp1 proteins were expressed in the *Hc* G217B cbp1 mutant background. We found that the chimeric proteins were expressed and secreted (Fig. 3D), suggesting that they were properly folded, but none could restore macrophage lytic capability (Fig. 3E). Interestingly, while *E. crescens* Cbp1 seems to block intracellular growth (Fig. 3C), chimeras of *E. crescens* Cbp1 with the Pb03 N-terminal residues necessary for lysis or *E. crescens* Cbp1 with the G217B C-terminal helical bundle no longer had this growth defect (Fig. 3F).

### *Es. africanus* Cbp1 is unable to cause macrophage lysis despite replicating intracellularly

To determine if the lack of lysis caused by *Emergomyces* Cbp1 homologs is reflective of the *Emergomyces* natural infection of primary macrophages, we infected BMDMs with *Es. africanus* wildtype yeast at an MOI of 1. Macrophage lysis was monitored by measuring the release of lactate dehydrogenase into the culture supernatants. We observed no lysis of infected macrophages over the course of infection (Fig. 4A). Tunicamycin treatment was used as a positive control for BMDM lysis (Fig. S4). Despite the lack of macrophage lysis, intracellular replication of *Es. africanus* was observed, and macrophages became filled with yeast cells over the course of the infection (Fig. 4B). This behavior is strikingly similar to how the *Hc cbp1* mutant grows intracellularly to high levels but cannot lyse host cells (33). The lack of macrophage lysis in *Hc* expressing the *Emergomyces* Cbp1 homologs is consistent with the inability of *Es. africanus* to lyse macrophages during infection. *Es. africanus* and other *Emergomyces* species are still capable of causing systemic disease in humans (39, 40) suggesting that these fungi may rely on virulence factors other than Cbp1.

**Fig. 4.**
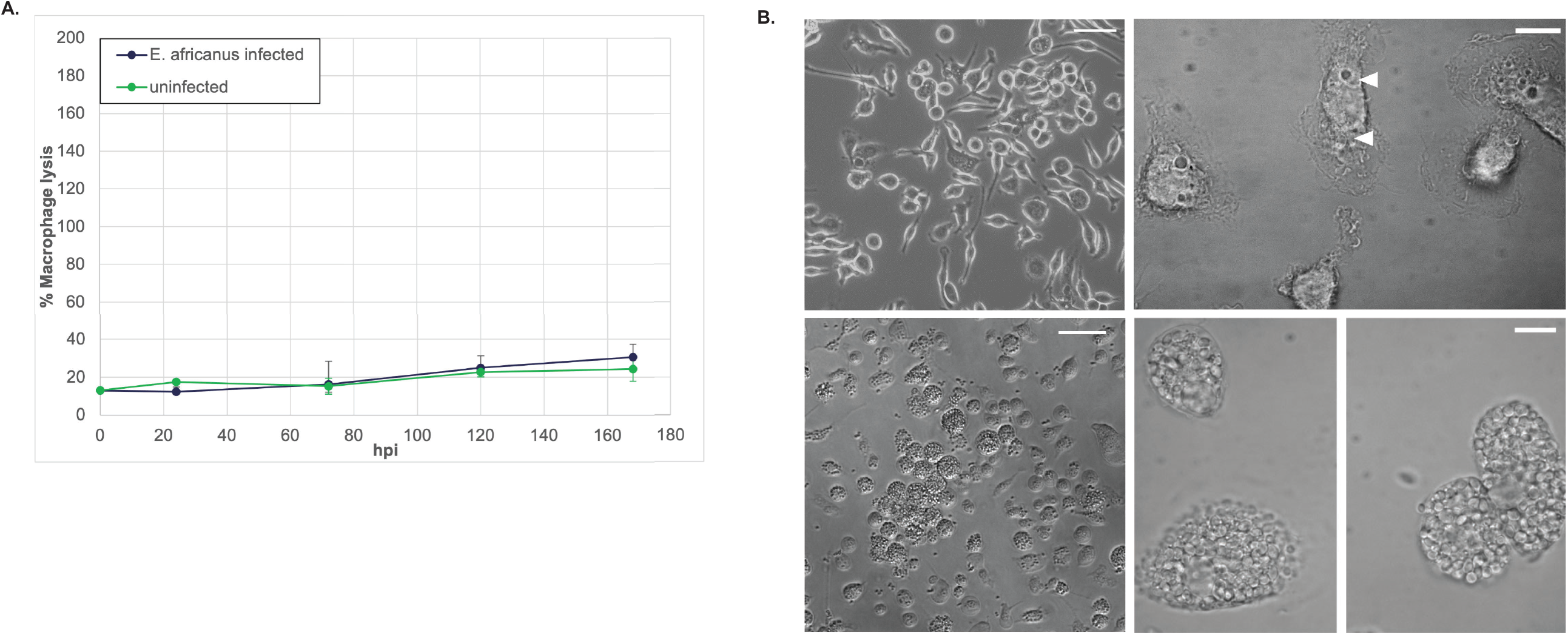
*Es. africanus* yeast can replicate intracellularly in macrophages but are unable to cause lysis. A. *Es. africanus* yeast were used to infect BMDMS at an MOI of 1 as described in the methods. LDH release was used to quantify percent host cell lysis. (B) DIC images of BMDMs infected with wildtype *Es. africanus* at an MOI of 1 at one day post-infection (top) or five days post infection (bottom), when multiple intracellular yeasts are present. Arrowheads indicate intracellular yeasts.

### *Hc* G217B and Pb03 Cbp1 enter the cytosol during macrophage infection

To further probe the mechanism of Cbp1 function, we investigated its subcellular localization during *Hc* infection. Once *Hc* is phagocytosed by macrophages, it begins to replicate intracellularly within a modified phagosomal compartment that fails to fuse with degradative lysosomes and maintains a neutral pH (26, 51, 52). We have previously shown that *Hc* infection triggers a cytosolic stress response inside of macrophages known as the Integrated Stress Response (ISR) that is dependent on Cbp1 (34). As in the case of *Hc* infection, if the cytosolic stress remains unresolved, the macrophage undergoes apoptotic cell death. Since Cbp1 seems to be a crucial secreted virulence factor and is required to cause a host cell cytosolic response, we hypothesized that Cbp1 gains access to the macrophage cytosol during infection. A Cbp1-GFP fusion was previously shown to localize to the *Hc*-containing phagosome (30), but localization of the native Cbp1 during infection has not been determined and it is unknown whether the Cbp1-GFP fusion is functional. Almost all attempts at tagging *Hc* Cbp1 render it non-functional, and peptide antibodies generated against Cbp1 recognize only the denatured form of the protein, precluding localization by indirect immunofluorescence. To overcome these limitations, we fractionated infected macrophages and monitored Cbp1 accumulation in the macrophage cytosol fraction vs. the membrane fraction, which includes the contents of intracellular vesicles. We separated the cytosolic fraction from the membrane fraction (validated by probing for alpha-tubulin, and calnexin respectively) and Cbp1 localized exclusively to the cytosolic fraction, suggesting it exits the *Hc*-containing phagosome during infection (Fig. 5A). To confirm that small endosomal compartments remained intact during the fractionation process, we determined that LAMP1 was not detectable in the cytosolic fraction (Fig. S5B). Additionally, we checked that the *Hc* yeast themselves are not rupturing during the fractionation protocol by confirming that the *Hc* transcription factor Ryp1 is present only in the fraction that contains whole *Hc* yeast, which are separated from the lysates along with the nuclear fraction prior to ultracentrifugation (Fig. S5A-B). Using a N-terminal FLAG-tagged Pb03 Cbp1 homolog that retains its lytic capability during infection (Fig. S6A), we confirmed the cytosolic localization of Pb03 Cbp1 via this subcellular fractionation approach (Fig. 5A). Interestingly, the analogous tagged version of *Hc* G217B Cbp1 is non-functional (Fig. S6B) but nonetheless localizes to the macrophage cytosol during infection, suggesting cytosolic localization alone is not sufficient to cause lysis of the host macrophage (Fig. 5A, Fig. S6). The functional *Pb03*-3XFLAG Cbp1 homolog appeared in a punctate pattern throughout the infected macrophage cytosol and occasionally overlapped with the *Hc*-containing phagosome, suggesting it could be accumulating and aggregating in the cytosol during infection (Fig. 5B).

**Fig. 5.**
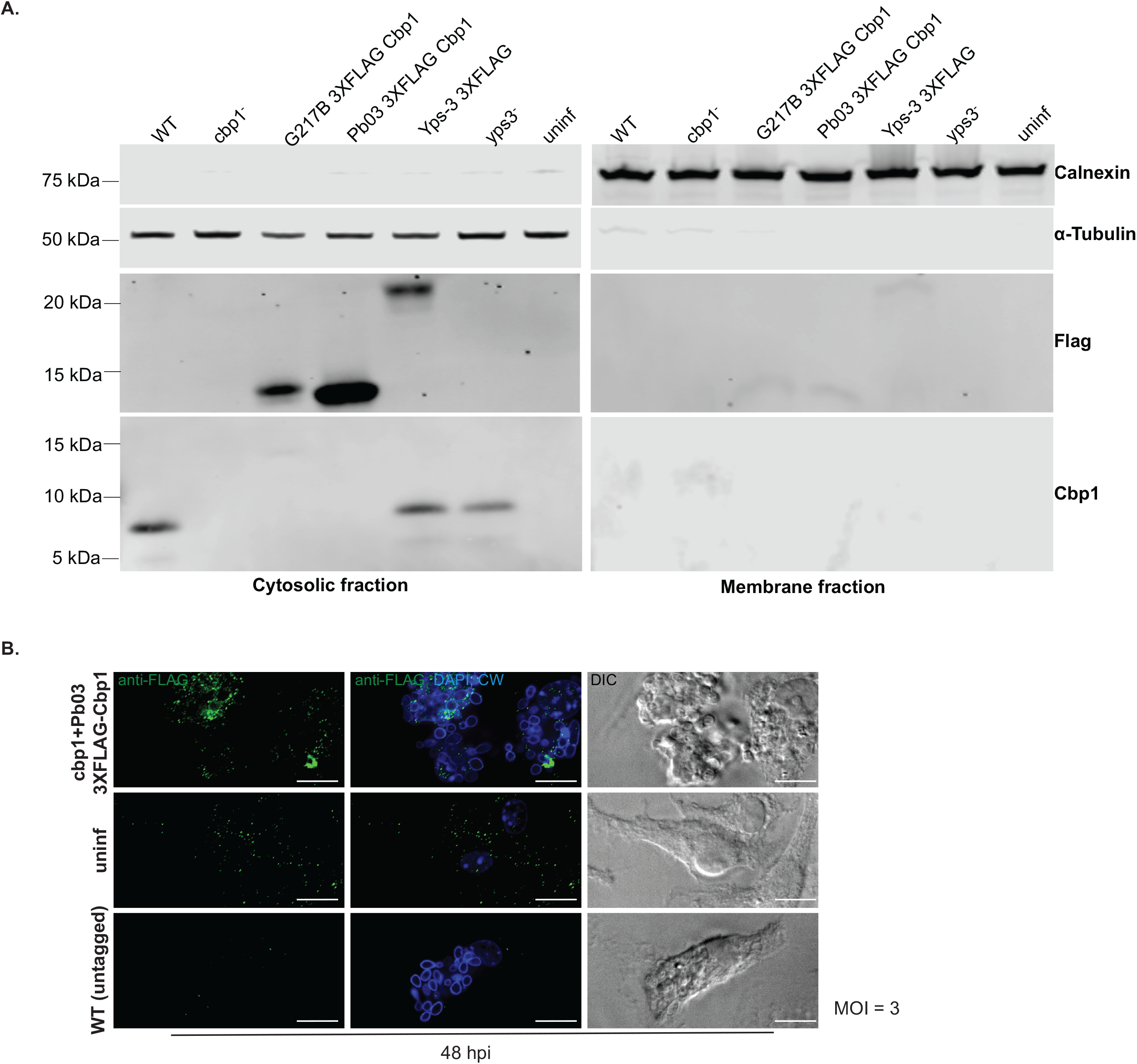
*Hc* and *Pb03* Cbp1 exit the *Hc-*containing phagosome and enter the macrophage cytosol during infection. A. BMDMs were mock-infected (uninf) or infected with either the “WT” strain (*Hc* G217B *ura5*^*-*^ carrying a *URA5* vector control), the *cbp1* mutant (*Hc* G217B *ura5*^*-*^ *cbp1*^*-*^) carrying either the vector control or G217B Cbp1 with 3XFLAG (G217B Cbp1 3XFLAG), Pb03 Cbp1 with 3XFLAG (Pb03 Cbp1 3XFLAG), WT G217B *ura5*^*-*^ carrying Yps-3 3XFLAG (Yps-3 3XFLAG), or the *Hc* G217B *ura5*^*-*^ *yps3*^*-*^ mutant (*yps3*^*-*^). Macrophage lysates were subjected to fractionation to separate cytosolic and membrane fractions, followed by SDS-PAGE and Western blotting using anti-Calnexin (marking the endoplasmic reticulum), anti-α-Tubulin (marking the cytosol), anti-FLAG or anti-Cbp1 antibodies. B. BMDMs were either mock-infected or infected with *Hc* G217B *ura5*^*-*^ *cbp1*^*-*^ expressing the WT untagged Cbp1(WT) or the *cbp1* mutant carrying the Pb03 Cbp1 tagged with 3XFLAG. Cells were fixed and subjected to indirect immunofluorescence with the FLAG antibody. Scale bar represents 10 μm.

### *Hc* Cbp1 forms a complex with Yps-3, another known *Hc* virulence factor that is also in the macrophage cytosol

To see if Cbp1 acts alone or as a part of complex of fungal proteins, we isolated 1xstrep-tagged G217B Cbp1 (Fig. S6C) from *Hc* culture supernatants and determined which *Hc* proteins associate with Cbp1 by mass spectrometry (Fig. 6A). The 1xstrep-tagged G217B Cbp1 allele retains partial lytic function during macrophage infection (Fig. S6C). In the eluates of the 1xstrep *Hc* G217B Cbp1 pulldown we observed a prominent band around 17-20 kDa that was absent in the control pull down of 2xstrep-eGFP. The identity of this band was later confirmed by mass spectrometry as *Hc* virulence factor Yps-3 (35-37). Conversely, Cbp1 co-purified with a pulldown of Yps-3-1xstrep from *Hc* culture supernatants (Fig. 6A). Additionally, our fractionation experiments demonstrated that, like Cbp1, Yps-3 is able to enter the macrophage cytosol during *Hc* infection (Fig. 5A).

**Fig. 6.**
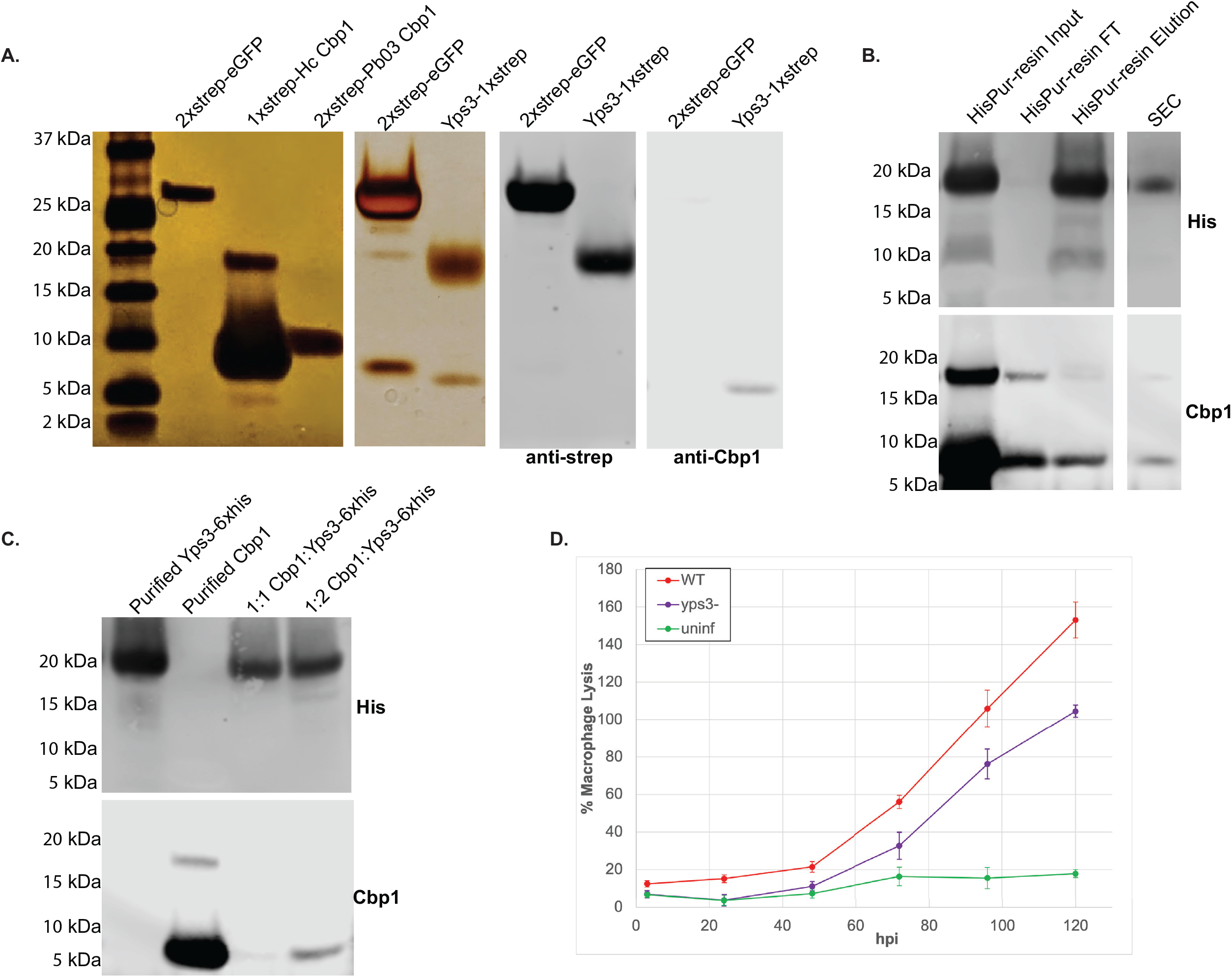
*Hc* G217B Cbp1 forms a complex with Yps-3. A. *Hc* culture supernatants from the *Hc* G217B *ura5*^*-*^ strain carrying a plasmid expressing 2xstrep enhanced GFP (2xstrep-eGFP) or Yps3-1xstrep, and the *Hc* G217B *cbp1* mutant expressing either G217B 1xstrep-Cbp1 or Pb03 2xstrep Cbp1 were subjected to StrepTactin affinity purification. Eluates were analyzed by SDS-PAGE followed by silver staining. The pulldowns of Yps-3-1xstrep and 2xstrep-eGFP were confirmed with SDS-PAGE followed with Western blot analysis with both anti-strep and anti-Cbp1 antibodies. B. *Hc* culture supernatant from the Yps-3-6XHis strain was subjected to SDS-PAGE followed by Western blotting with either anti-His or anti-Cbp1 antibodies. The left-hand panel shows input, flow-through (FT) and elution from the HisPur resin whereas the right-hand panel shows the fractions containing both Yps3 and Cbp1 from a size exclusion column (SEC). C. Purified Yps3-6XHis and purified G217B Cbp1 were either subjected separately to SDS-PAGE and Western blotting (lanes 1 and 2) or mixed first at defined molar ratios and then isolated by Ni-NTA pulldown of Yps-3-6XHis (lanes 3 and 4). D. BMDMs were either mock-infected (uninf) or infected at an MOI = 1 with G217B *ura5-* (WT) or G217B *yps-3*Δ, each transformed with a *URA5* vector. LDH release was calculated at multiple timepoints post-infection.

To determine if Yps-3 and Cbp1 form a complex, we purified C-terminally tagged Yps3-6xhis from culture supernatants using a cobalt resin and observed that Cbp1 co-purifies with Yps-3 (Fig. 6B). To isolate pure Yps3-6xhis, we used a cation exchange column, but a large portion of Yps-3 was present in the flow through of the cation exchange column along with Cbp1. To confirm that the proteins are truly in a complex, we passed the flow through over a sizing column and discovered that Yps-3 and Cbp1 were present in the same fractions, suggesting they remained in a tight complex (Fig. 6B). Additionally, we mixed purified Yps3-6xhis and G217B Cbp1 in molar ratios of 1:1, 1:2, 2:1, 5:1 and isolated the 6xHis tagged Yps3 with a Ni-NTA resin. We found that the molar ratio of 1:2 Cbp1:Yps-3 yielded the most robust interaction between the two proteins (Fig. 6C, Fig. S7). Previously Yps-3 RNAi knockdown had shown no yeast phenotype in a macrophage cell line (35). Since

RNAi may not be definitive, to further characterize the role of Yps-3 in macrophage infection we generated a CRISPR deletion mutant of *YPS-3* (Fig. S8A and B) (53, 54). We found that the *yps-3*Δ mutant has a partial lysis defect in BMDMs (Fig. 6D) despite the presence of Cbp1 in the cytosol of macrophages infected with this mutant (Fig. 5A). These data indicate that Yps-3 is required for robust lysis of infected macrophages, perhaps because it potentiates the activity of Cbp1.

## Discussion

Macrophages are innate immune cells that are critical for the early detection and killing of infectious microbes. *Hc* is an intracellular pathogen that evades macrophage anti-microbial mechanisms, thereby surviving and replicating within a modified phagosomal compartment. Here we focus on a secreted *Hc* effector, Cbp1, that is critical for manipulating the macrophage response. Cbp1 is absolutely required for macrophage lysis during *Hc* infection. Using cutting-edge bioinformatics tools, we identified all existing Cbp1 homologs in the fungal kingdom and interrogated these proteins for their ability to trigger macrophage lysis when expressed in a *Hc cbp1* mutant. These experiments revealed that only *Hc* and *Pb* carry “lytic” Cbp1 alleles, whereas closely related *Emergomyces* Cbp1 variants cannot promote macrophage lysis during *Hc* infection. Interestingly, the crystal structure of *Hc* and *Pb* Cbp1 proteins uncovered a novel fold in this rapidly evolving lysis factor.

Our previous work revealed that *Hc* Cbp1 is absolutely required for triggering an Integrated Stress Response (ISR) in the macrophage cytosol during *Hc* infection. When we mutagenized Cbp1 to yield alleles that were partially lytic or completely defective for lysis, we observed little or no ISR induction, respectively, indicating a correlation between the ability of Cbp1 to trigger the ISR and lyse macrophages (34). Here we show for the first time that *Hc* Cbp1 accesses the macrophage cytosol during infection, suggesting that the site of action of Cbp1 is in the cytosol of the host cell. Additionally, we show that Cbp1 binds another known *Hc* virulence factor, Yps-3. Previous work in the literature showed that Yps-3 localizes to the *Hc* cell wall and is released into culture supernatant. RNA interference strains targeting Yps-3 were used to show a role for Yps-3 in organ colonization in the mouse model of infection but no defect in macrophage lysis was observed at the single timepoint examined (35). Interestingly, we show here that deletion of Yps-3 results in diminished host-cell lysis, suggesting that Yps-3 could help potentiate the function of Cbp1.

How Cbp1 and Yps-3 effectors transit from the phagosome to the macrophage cytosol is not yet clear. Perhaps the *Hc-*containing phagosome is leaky, thereby allowing small fungal proteins to access the cytosol. Alternatively, *Hc* is known to secrete copious amounts of extracellular vesicles that contain varied cargo of *Hc* proteins (55, 56).These virulence factors could enter the cytosol because exosomes, or even multi-vesicular bodies, fuse with the phagosome membrane and release their contents directly into the cytosol. Once in the reducing environment of the host cytosol, perhaps Cbp1 changes its conformation and triggers the ISR. We observed that a tagged allele of the Pb03 Cbp1 (Pb03*-*3XFLAG Cbp1) was distributed throughout the cytosol in a punctate pattern. We speculate that Cbp1 could be aggregating in the cytosol and triggering cellular stress, and future work will focus on the mechanism of action of the Cbp1-Yps-3 complex in triggering host-cell death.

We observed that Cbp1 appears to be a rapidly evolving lysis factor that is undergoing positive selection. Since Cbp1 is present in all of the genomes of the Ajellomycetaceae human fungal pathogens except *Blastomyces* species, it is likely that Cbp1 arose at the base of the Ajellomycetaceae tree and was subsequently lost only in *Blastomyces*. Notably, *Blastomyces* yeast cells are largely extracellular during infection of mammals, in contrast to *Histoplasma* and other Ajellomycetaceae, suggesting that Cbp1 could be an adaptation to an intracellular lifecycle. Interestingly, despite its lack of Cbp1, *Blastomyces* has retained a Yps-3 homolog, *Blastomyces* Adhesin 1 (Bad1). Bad1 mediates binding of *Blastomyces* to host cells and plays an immunoregulatory role during infection (42, 57-60). In the case of *Hc*, it is possible that Yps-3 acts with Cbp1 to promote macrophage lysis as well as playing a Cbp1-independent role during infection.

*Hc* Cbp1 is critical for macrophage lysis and dissemination (33, 34). Pb18 Cbp1 is highly expressed in *Paracoccidioides* Pb18 yeast cells (61) and Pb03 Cbp1 is capable of triggering host-cell lysis when expressed in the *Hc* G217B *cbp1* mutant background, but whether any *Paracoccidioides* species utilize Cbp1 during infection is yet to be determined. In contrast, *Emergomyces* species have Cbp1 in their genomes but heterologous expression of these Cbp1 alleles in *Hc* suggests that they are non-lytic. This observation correlates with the natural infection of primary BMDMs by *Es. africanus*, characterized here for the first time. We observed that *Es. africanus* undergoes intracellular replication within macrophages but is unable to lyse them, much like the *Hc* G217B *cbp1* mutant. It is unclear whether the absence of a lytic Cbp1 affects *Emergomyces* pathogenesis. Notably *Emergomyces* species are pathogens of immunocompromised people (62). whereas *Hc* is a primary pathogen that can infect healthy individuals, indicating that *Hc* may harbor more potent virulence strategies. Nonetheless, *Emergomyces* species can disseminate and cause disease in their hosts despite the likelihood that their Cbp1 proteins are non-lytic.

The analysis of *Hc* H88 and Pb03 Cbp1 protein structures revealed a novel fold that was highly distinct from the previously published NMR G186AR Cbp1 structure. Our protein purification strategy was non-denaturing, which contrasts to the methodology used for the NMR structure. It is possible that Cbp1 can fold differently depending on its environment; for example, perhaps the reducing environment of the macrophage cytosol triggers a conformational change in Cbp1 protein. When we compared the two new structures of Pb03 and H88 Cbp1, we found that charge was not distributed similarly across the proteins. The bottom of the C-terminal helices that form the core four-helix bundle of the Pb03 Cbp1 are much more negative relative to the *Hc* H88 Cbp1 homolog. Interestingly, only the Pb03 (and not H88) structure shows partially negative N-terminal helices. In addition, when we overlaid the previously published alanine scan of G217B Cbp1 over the two new structures, we found that the majority of the residues necessary for optimal lysis are distributed in the N-terminal helix, suggesting that this region of the protein mediates host-cell death. However, when we swapped the 1^st^ helix of the non-lytic *E. crescens* Cbp1 allele with the corresponding helix from either G217B or Pb03 Cbp1, the resulting chimeric protein was not lytic, suggesting that other regions of the protein may affect folding and/or function. Nonetheless, the concordance between the alanine scan phenotypes and the location of the residues in the crystal structure gives strong support to our analysis of the Cbp1 fold and indicates that our structural work on Cbp1 is a critical step in understanding how intracellular fungal pathogens manipulate host cell viability.

## Materials and Methods

### Primers

Primers used in this study are described in Table 2.

### Determining the conservation of Cbp1 in Ajellomycetaceae genomes and generating protein alignments

The Cbp1 protein alignment from Fig S3 of (34) was extended as follows: First, we used HMMer 3.1b2 (63) to build an HMM from the previous alignment and used it to search for homologs in the following annotated protein sets, downloaded from Genbank: *Emergomyces africanus* (GCA_001660665.1), *Emergomyces pasteuriana* (GCA_001883825.1), *Emmonsia parva* (GCA_001014755.1), *Blastomyces percursus* (GCA_001883805.1), *Paracoccidioides venezuelensis* Pb300 (GCA_001713645.1) and Paracoccidioides restrepiensis *PbCNH* (GCA_001713695.1). This yielded single homologs in *Es. africanus, Pb300*, and *PbCNH*, two homologs in *Es. pasteuriana*, and no homologs in *E. parva* or *B. percursus*, which is consistent with loss of *CBP1* in genus Blastomyces. The exon annotations of the five new homologs were refined based on the previous protein alignment. Four of the new homologs are syntenic to the previously known Cbp1s and share the conserved two intron structure. The remaining homolog, from *Es. pasteuriana*, is in a distinct genomic location and has three introns. TBLASTN from NCBI BLAST 2.6.0 (64) was then used to search the full set of Cbp1 protein sequences against the unannotated *Emergomyces orientalis* genome (GCA_002110485.1), yielding one hit orthologous to the conserved two intron Cbp1 sequence and one hit orthologous to the three intron *Es. pasteuriana* paralog. Exons were annotated for both *Es. orientalis* homologs based on the existing protein alignment, as above. All sequences were then aligned with PROBCONS 1.12 (65) to yield the final alignment.

### Species tree generation

Full length midasin sequences were inferred by using NCBI TBLASTN 2.6.0 (64) to search the G217B predicted sequence HISTO_DA.Contig93.Fgenesh_Aspergillus. 103.final_new against each genome of interest, retaining the top non-redundant hits spanning the full protein. An HMM was built from the initial TBLASTN alignments with hmmbuild (HMMer 3.1b2) (63), all hits were realigned to the HMM with hmmalign (HMMer 3.1b2), and the resulting multiple alignment was refined with PROBCONS 1.12 (65). A phylogenetic tree was inferred from the ungapped positions of the PROBCONS multiple alignment with IQTREE 1.5.3(66).

### PAML analysis of positive selection of Cbp1

The full length CBP1 protein alignment was cut to the 11 CBP1 orthologs (dropping the paralogs from *Es. pasteuriana* and *Es. orientalis*) and was converted to a nucleotide alignment by replacing amino acids with their cognate codons from the respective genome sequences. The corresponding evolutionary tree was taken from the midasin-based phylogeny, cutting the tree to just species with CBP1 and dropping branch lengths. CODEML from PAML 4.9(44) was used to fit the nucleotide alignment and tree topology to four evolutionary models (M1a: neutral, M2a: positive selection, M7: beta, and M8: beta and omega). Models with and without positive selection were then compared by using CHI2 from PAML 4.9 to perform chi squared tests on the corresponding log likelihood ratios from CODEML with two degrees of freedom. The corresponding p-values are reported in the text. Specific positions under positive selection with a posterior probability greater than 0.95 based on model M2a were identified using the Naive Empirical Bayes output from CODEML.

### BMDM culture conditions

Bone marrow derived macrophages (BMDMs) were isolated from the tibias and femurs of 6-8 week old C57BL/6J (Jackson Laboratories stock no. 000664) mice. Mice were euthanized via CO2 narcosis and cervical dislocation as approved under UCSF Institutional Animal Care and Use Committee protocol # AN181753-02B. Cells were differentiated in BMM (bone marrow derived macrophage media) which consists of Dulbecco’s Modified Eagle Medium, D-MEM High Glucose (UCSF Cell Culture Facility), 20% Fetal Bovine Serum (Atlanta Cat #: S11150, Lot #: D18043), 10% v/v CMG supernatant (the source of CSF-1), 2 mM glutamine (UCSF Cell Culture Facility), 110 μg/mL sodium pyruvate (UCSF Cell Culture Facility), penicillin and streptomycin (UCSF Cell Culture Facility) with 20% Fetal Bovine Serum (Atlanta Cat #: S11150, Lot #: D18043). Undifferentiated monocytes were plated in BMM that contains BMM for 7 days at 37°C and 5% CO_2_. Adherent cells were then scraped and frozen down in 40% FBS and 10% DMSO until further use. *Hc* cultures were grown in liquid *Histoplasma* macrophage media (HMM) using an orbital shaker or on HMM agarose plates (33).

### Generation of Hc strains

*H. capsulatum* strain G217B *ura5Δ* (WU15) was a kind gift from the lab of William Goldman (University of North Carolina, Chapel Hill). For all studies involving the *cbp1* and *yps-3* mutants, “wildtype” refers to G217B *ura5Δ* transformed with a *URA5*-containing episomal vector (pLH211), *cbp1* refers to G127Bura5 *Δcbp1::T-DNA* as previously described (33) transformed with the same *URA5*-containing episomal vector, and “complemented” strain refers to *G217Bura5Δcbp1::T-DNA* transformed with the *URA5*-containing plasmid bearing the wild-type CBP1 gene (pDTI22) as previously described (33).The *Emergomyces CBP1* coding sequences (*Es. africanus, Es. orientalis, E. crescens, Es. pasteuriana_1, Es. pasteuriana_2)* and *E. crescens> G217B* or *E*.*crescens>Pb03* chimeric constructs (N-terminal helix residues, C-terminal loop residues, both C-terminal and N-terminal residues, 1^st^ helix swap) constructs were synthesized as gBlocks™ by Integrated DNA Technologies and cloned into pDTI22, replacing the G217B *CBP1* coding sequence but maintaining the flanking sequences. *P. americana* strain *Pb03* Cbp1 gene construct includes the *Hc* G217B Cbp1 signal peptide fused to the mature Pb03 protein coding sequence flanked by the same regulatory sequences of pDTI22 as previously described (34). All tagged Cbp1 expressing strains including 1xstrep-H*c G217B* Cbp1, 2xstrep-*Pb03* Cbp1, 3XFLAG-*Hc G217B* Cbp1, 3XFLAG-*Hc Pb03* Cbp1, were N-terminally tagged with the tag situated between the G217B Cbp1 signal peptide and the mature protein sequence and were flanked by the same regulatory sequences in pDTI22 and introduced into the Hc G217B cbp1 mutant strain. The *yps3* mutant was generated from the G217B *ura5Δ* parental strain transformed with the episomal plasmid pDAZ021 which contains a bidirectional H2Ab promoter driving Cas9 and two sgRNA cassettes targeting the sequences on both sides of the G217B Yps-3 gene (54). Single colony isolates for PCR screened in the yps-3 genomic locus to look for excisions in the colony population and the isolates with the most likely edited band were struck out for further single colony isolation until an isolate with a clean excision between the two sgRNA target sites was detected. pBJ219 was subsequently lost from the mutant by growing the *Hc* yeast in the presence of exogenous uracil and then screening for plasmid loss. Subsequently, the G217B *ura5Δ yps3*Δ was transformed with the *URA5-*containing episomal plasmid (pLH211). The overexpression strains used to detect and purify Yps-3 were generated by transforming G217B *ura5Δ* parental strain with an episomal plasmid with each of the C-terminally tagged Yps-3 alleles (Yps-3-6xHistidine, Yps-3-2xstrep, Yps-3-3XFLAG) flanked by the Cbp1 promoter and CatB terminator. The control for the StrepTactin pulldowns was an N-terminally Twinstrep™ (iba Lifesciences) tagged enhanced GFP (eGFP) that was fused to the G217B Cbp1 signal peptide and cloned into the same episomal plasmid (pDTI22) as all the other Cbp1 constructs. For all Cbp1, Yps3, eGFP, and CRISPR construct plasmids, approximately 50 ng of PacI-linearized DNA was electroporated into the appropriate parental strain (G217B *ura5Δ*, G217B ura5*Δ cbp1::T-DNA*, or G217B ura5*Δ yps3*Δ) as previously described (33). The plasmid pLH211 was used as a vector control. Transformants were selected on HMM agarose plates.

### Hc secreted protein detection in culture supernatants

To detect *Hc* secreted proteins (tagged Cbp1 alleles and Cbp1 homologs, tagged Yps-3 homologs, tagged 2xstrep-eGFP) in culture supernatants, 4-5 day *Hc* cultures were grown in liquid HMM and yeast were pelleted by centrifugation. The supernatants were subjected to filtration using 0.22 μm filters, and the resultant filtrates were concentrated using Amicon Ultra Centrifugal Filter Units with a 3 kDa cutoff (EMD Millipore). Protein concentration was quantified using the Bio-Rad protein assay (Bio-Rad Laboratories). Equal amounts of protein were separated by SDS-PAGE, and proteins were visualized by staining the gel with InstantBlue™ Coomassie Protein Stain (ISB1L – abcam 119211)

### Macrophage Infections

Macrophage infections with G217B *Hc* strains were performed as described previously (33). Briefly, the day before infection, macrophages were seeded in tissue culture-treated dishes. On the day of infection, yeast cells from logarithmic-phase *Hc* cultures (OD_600_ = 5–7) were collected, resuspended in BMM, sonicated for 3 seconds on setting 2 using a Fisher Scientific Sonic Dismembrator Model 100, and counted using a hemacytometer at 40X magnification. Depending on the multiplicity of infection (MOI), the appropriate number of yeast cells was then added to the macrophages. After a 2-hour phagocytosis period, the macrophages were washed once with d-PBS to remove extracellular yeast and then fresh media was added. For infections lasting longer than 2 days, fresh media was added to the cells at approximately 48 hours post infection.

### Fractionation of Hc-infected macrophage lysates

2.5×10^7^ BMDMs were plated on a TC-treated 15-cm plates and allowed to adhere for at least 24 hrs. The BMDMs were then infected with *Hc* at an MOI = 5 and the infection was allowed to proceed for 24 hours. At 24 hours post infection, cells were collected by scraping without washing and spun down at 2500 rpm for 5 min to pellet the intact cells. The cell pellet was then resuspended in 500 μL of d-PBS, transferred to a 1.5 mL tube, and spun at 1000 rpm for 2 min. The cell pellet was then resuspended in 300 μL of homogenization buffer (150 mM KCl, 20 mM HEPES pH 7.4, 2 mM EDTA, cOmplete Mini Protease Inhibitor Cocktail tablet-Roche 04693124001). For the unfractionated sample, the cell pellet was resuspended in homogenization buffer with 1% TritonX-100. For the fractionated samples, the cell lysate was then gently passaged through a 27-gauge needle to disrupt only the plasma membrane and not any internal membranous compartments. The lysate was then spun at 3000 rpm for 5 minutes to pellet *Hc* cells and macrophage nuclei. The remaining lysate was then further clarified by centrifuging twice at 3000 rpm with removal of 275μL and 250 μL respectively to avoid contamination of the lysate with *Hc*. After the final low speed spin, 225 μL was placed in ultracentrifuge tubes (Beckman Coulter 349622), weighed, and loaded into a Beckman Coulter Fixed Angle Rotor TLA100.3. The samples were then spun at 45,000 rpm for 2 hours in a TL-100 tabletop ultracentrifuge to pellet the membranes. After the spin, 175 μL of supernatant, representing the cytosolic fraction, was transferred into a separate 1.5 mL tube. The remaining supernatant was discarded to prevent cross-contamination between the cytosolic and membrane fractions. The pellet, representing the membrane fraction, was resuspended in 225 μL of homogenization buffer with 1% TritonX-100. All samples were flash frozen in liquid nitrogen and stored at -80°C.

### Lactate dehydrogenase release assay

To quantify macrophage lysis, BMDMs were seeded (7.5 × 10^4^ cells per well of a 48-well plate) and infected as described above. At the indicated time points, the amount of LDH in the supernatant was measured as described previously (33). BMDM lysis was calculated as the percentage of total LDH from supernatant of the uninfected wells with uninfected macrophages lysed in 1% Triton X-100 at the time of infection. Due to continued replication of BMDMs over the course of the experiment, the total LDH at later time points is greater than the total LDH from the initial time point, resulting in an apparent lysis that is greater than 100%.

### Intracellular replication

BMDMs were seeded (7.5×10^4^ cells per well of a 48-well plate) and infected in triplicate as described above. At the indicated timepoints, culture supernatants were removed and 500 μl of ddH2O was added. After incubating at 37°C for 15 min, the macrophages were mechanically lysed by vigorous pipetting. The lysate was collected, sonicated to disperse any clumps, counted, and plated on HMM agarose in appropriate dilutions. After incubation at 37°C with 5% CO_2_ for 12–14 days, CFUs were enumerated. To prevent any extracellular replication from confounding the results, intracellular replication was not monitored after the onset of macrophage lysis.

### Purification, crystallization and structural determination of Cbp1

2-3 L of G217B *Hc* expressing each different *cbp1* allele were grown at 37°C with 5% CO_2_ for 5 days. Culture medium was concentrated using an Amicon cell outfitted with a 5kDa molecular weight cutoff membrane. Concentrated medium was diluted 1:50 in MonoA buffer (20 mM Tris-HCl pH 7.5, 20 mM NaCl) and run through a HiTrapQ ion exchange column. Protein generally eluted at a 24% MonoB buffer concentration (20 mM Tris-HCl pH 7.5, 1 M NaCl). Eluted fractions were pooled and concentrated using a 3 kDa molecular cutoff spin concentrator. The final product was separated from remaining contaminants using a size exclusion approach with a Superdex 75 10/300 column run with SEC buffer (50 mM Tris-HCl pH 7.5, 150 mM NaCl). Purity of the final elution was verified by SDS-PAGE and concentrated using a 3 kDa MWCO spin concentrator. Pb03 Cbp1 was concentrated to 6 mg/mL and H88 Cbp1 was concentrated to 14 mg/mL. Crystallization trays were set up in sitting drop vapor diffusion trays at room temperature using 2 μL + 2 μL drops of protein and crystallization solution. Crystals of Pb03 Cbp1 formed in 0.05 M HEPES pH 6.5, 35% PEG 6000, and were seen after 24 hours and allowed to grow for 48 hours to reach full size. H88 Cbp1 crystallized in 0.02 M CoCl2, 0.2 M MES pH 6.5, 2 M Ammonium sulfate using the same setup as for Pb03 Cbp1, crystals were observed after 48 hours and reached full size in 1 week. To provide phase information, full sized Pb03 Cbp1 crystals were soaked for 2 hours in 10 mM potassium tetrachloroplatinate (II) from the Hampton Heavy Atom kit screen. Derivatized and native Pb03 Cbp1 crystals were cryoprotected using ethylene glycol. H88 Cbp1 crystals were cryoprotected using glycerol. All diffraction data were collected at ALS BL 8.3.1 on a Pilatus3 S 6M detector. Data processing and refinement was conducted in the CCP4 and Phenix programming suites. Datasets were processed using XDS and sailed using AIMLESS in the CCP4 suite. The structures were solved using Phaser in the Phenix crystallography suite.

### BMDM infection with Emergomyces africanus yeast

*Emergomyces africanus* (clinical isolate CBS 136260 (67, 68)), was grown in brain-heart infusion broth (BHI) (Cat No. 110493, Merck, USA), at 37 °C, 180 rpm, for 5 days. Yeasts were then sub-cultured for a further 3 days by 1:20 dilution into fresh BHI, until reaching an optical density of 3-4. BMDMs were seeded in 8-well μ-Slide imaging slides (Cat. No. 80826, ibidi, Germany) at a density of 7.5 × 10^4^ cells/well in 0.375 ml BMM (DMEM, high glucose, GlutaMAX™ supplement, pyruvate (Thermo Fisher, Cat no. 10569010), 20% FBS (Thermo Fisher, Cat. no. 10270106), Penicillin-streptomycin (50 units/ml) (Thermo Fisher, Cat. no. 15140148), 20 ng/ml mCSF (R&D Systems, Cat no. 416-ML)), and incubated at 37 °C, 5% CO_2_ for 24 h. Yeast cells used for infection were washed twice by centrifugation and resuspension in PBS, counted using a hemocytometer, then the appropriate volume of yeast suspension was added to BMM. On day 0, BMDMs were infected by replacing the media with 0.375 ml yeast suspension in BMM, with a total of 7.5 × 10^4^ yeasts/well (MOI=1). The BMDMs were fed on day 2 by removing 200 μl spent media from wells and replacing it with 200 μl fresh BMM.

### LDH assay sample collection of BMDMs infected with Es. africanus

BMDMs were seeded in 48-well tissue culture plates at a density of 1 × 10^5^ cells/well in 0.5 ml BMM, and incubated at 37 °C, 5% CO_2_ for 24 h. Yeast cells used for infection were washed twice by centrifugation and resuspension in PBS, counted using a hemocytometer, then the appropriate volume of yeast suspension was added to BMM. On day 0, BMDMs were infected by replacing the media with 0.5 ml yeast suspension in BMM, with a total of 1 × 10^5^ yeasts/well (MOI=1). Media was replaced with 0.5 ml fresh BMM for wells containing uninfected BMDMs. Uninfected BMDMs were lysed on day 0 by removing the media and replacing it with 0.5 ml 0.2 % Tween-20 in distilled water, and the lysate was collected and stored in 1.5 ml microcentrifuge tubes at 4 °C. The BMDMs were fed on day 2 by adding 200 μl BMM. On days 0, 1, 3, 5 and 7, spent media (175 μl) was collected from wells in and stored in 1.5 ml microcentrifuge tubes at 4 °C for later analysis in the LDH assay.

### Imaging of Es. africanus infected BMDMS

On imaging days, the media was removed and replaced with 250 μl pre-warmed 4% PFA for 30 min. The PFA was then removed and replaced with PBS. The slides were imaged using a Zeiss Axiovert 200M inverted fluorescence microscope with a Zeiss AxioCam HSc camera.

### Immunofluorescence of FLAG-Pb03 Cbp1 during macrophage infection

1.5×10^5^ BMDMs were plated on 12 mm circular coverslips (Fisher Scientific 12-545-80) in a 24-well plate and infected as described above with the following *Hc* strains at an MOI of 3; cbp1 mutant with wildtype G217B Cbp1 and 3XFLAG-Pb03 Cbp1. At 24 and 48 hours post infection cells were fixed with 4% Paraformaldehyde for 15 minutes at room temperature and washed 3 times with d-PBS to remove residual paraformaldehyde. Fixed cells were permeabilized and nonspecific binding sites were blocked with a filter-sterilized Saponin Block solution (0.2 % Saponin from Quillaja Bark (Sigma-Aldrich S7900), 2% heat inactivated FBS (Atlanta Cat #: S11150, Lot #: D18043)) for 1 hour at room temperature. Cells were stained a monoclonal primary mouse anti-FLAG M2 antibody (Millipore Sigma F3165) at a dilution of 1:250 and incubated in a light-blocking incubation chamber overnight with 4°C. Coverslips were washed 3 times with block for 5 min to wash off any remaining primary antibody. Coverslips were incubated with a polyclonal **I**gG (H+L) Highly Cross-Adsorbed Goat anti-Mouse, Alexa Fluor™ 488 (Fisher Scientific A11029) secondary at a dilution of 1 drop into 500 μL of block for 1 hour at room temperature in a light-blocking incubation chamber. Coverslips were then washed 3 times with d-PBS for 5 minutes and then briefly rinsed in double distilled water and the excess liquid was absorbed by the corner of a kimwipe. Coverslips are mounted on glass slides with a drop of Vectashield Mounting media with DAPI that has 1:100 Calcofluor White (1 mg/mL). Coverslips were sealed to the glass slide with clear nail polish. Images were taken on a Nikon CSU-X1 spinning disk confocal microscope

### Affinity purification of 1x/2xstrep-tagged secreted proteins

1xstrep/2xstrep-tagged proteins (2xstrep-eGFP, 1xstrep G217B *Hc* Cbp1, 2xstrep *Pb03* Cbp1, and Yps-3-1x/2xstrep) were purified from either *Hc* culture supernatants or from *Hc*-infected macrophage lysates. In the case of *Hc* culture supernatants, 25 mL of *Hc* culture was grown for 4 days at 37°C with 5% CO_2_ and passed through a 0.22 μm syringe filter. Roche cOmplete Mini Protease Inhibitor Cocktail tablet (Roche 04693124001) was added to the filtered supernatant which was then concentrated using 3000 MWCO Amicon 15 mL centrifugal filters (Millipore Sigma UFC900324) to approximately 500 μL and then diluted back to 1 mL total with d-PBS. To generate *Hc*-infected macrophage lysates, 4 plates of BMDMs seeded at a density of 2.25×10^7^ were infected with *Hc* at an MOI = 5. At 24 hours post infection, the adherent cells were scraped in 10 mL of cold d-PBS and pelleted at 1500 rom for 10 min at 4°C. After decanting the d-PBS, the cell pellet was suspended in 1 mL of cold lysis buffer (50 mM TRIS-HCl pH 7.5, 150 mM NaCl, 1 mM EDTA, 0.5% NP-40, 1 tablet of Roche cOmplete Protease inhibitor Cocktail, and 1 tablet of Millipore Sigma PhosStop (4906845001). The lysate was sonicated 2 times for 5 seconds at setting 2 using a Fisher Scientific Sonic Dismembrator Model 100 with 1 min on ice in between. The cells were further lysed for 1 hr at 4°C with a rotisserie revolver and the lysate was clarified with a 20 min spin at 13,000 rpm at 4°C. Both the *Hc* supernatant and macrophage lysates were applied to pre-washed Magstrep “type3” XT beads (iba Life Sciences 2-4090-002) and allowed to bind overnight at 4°C on a rotisserie revolver. The Magstrep beads with bound strep-tagged proteins were then washed three times with wash buffer Buffer W (iba Lifesciences 2-1003-100) and eluted with Buffer BXT (iba Lifesciences 2-1042-025).

### Yps3-6xHis purification

3 Liters of G217B *Hc* expressing Yps3-6xHis were grown at 37°C with 5% CO_2_ for 5 days. The supernatant was sterile filtered through a 0.22 μm 0.5 or 1L vacuum filter and concentrated using the Amicon 200 mL Stirred Cell (UFSC20001) with a 3 kDa NMW Ultrafiltration Disc (Millipore PLBC06210) and Amicon Ultra 15 mL Centrifugal Filters with a 3kDa NMW (Millipore UFC9003). The concentrated supernatant was applied to a cobalt resin to achieve a His-tag pulldown using the HisPur™ Cobalt Purification Kit (ThermoScientific™ 90092). The elutions were then applied to a cation exchange column to purify Yps3-6xhis. The majority of the Yps-3-6xhis emerged with Cbp1 in the flow-through.

### Yps3-6xhis and Cbp1 protein interaction assays

Concentrated supernatant was diluted 1:50 in Mono A buffer (20mM Tris-HCl pH 7.0, 20 mM NaCl). The sample was applied to a HitrapS cation exchange column and eluted with 25% MonoB buffer (20 mM Tris-HCL pH 7.0, 1 M NaCl). The flow-through fractions contained both Cbp1 and Yps3-6xHis, as a complex of these proteins would retain a negative change at the buffered pH. The complex was validated by passage through a Superdex75 Size Exclusion column using 50 mM Tris pH 7.0, 150 mM NaCl. Both Cbp1 and Yps-3-6xHis eluted in a monodispersed peak. To determine at what molar ratios this complex of Yps-3 and Cbp1 forms at pure Yps-3-6xHis and G217B Cbp1, were mixed in different molar ratios and pulled down with NiNTA resin. For the Yps3 - 6xhis pulldown, the SEC flowthrough that contained both Cbp1 and Yps3-6xhis was incubated with 20 uL of washed 50% NiNTA bead slurry for 30 minutes at room temperature with shaking and then the beads were washed with 20 column volumes of NiNTA wash buffer (50 mM Tris-HCl pH 7, 150 mM NaCl, and 30 mM of Imidazole). The complex was eluted with 6 column volumes of NiNTA elution buffer (50 mM Tris-HCl pH 7, 150 mM NaCl, 500 mM of Imidazole). We verified the identity of the proteins in the complex by western blot using native antibody for Cbp1 and a mouse anti-his tag antibody (ABclonal AE003) for Yps3-6xhis.

### Mass Spectrometry

This work used the Vincent J. Proteomics/Mass Spectrometry Laboratory at UC Berkeley, supported in part by NIH S10 Instrumentation Grant S10RR025622. For gel band, 1D, and 2D MUDPIT Mass spectrometric analysis, trypsin or chymotrypsin were used to digest protein into peptides according to Facility protocols.

### SDS-PAGE Protein Gel and Western Blot analysis

Protein samples were mixed with 4X Protein Loading Buffer (Li-COR 926-31097) and 1:20 DTT (1M DTT) and denatured at 95°C for 5 minutes. Proteins were separated by SDS-PAGE on NOVEX-NuPAGE 4-12% BIS-TRIS gels with MES Running Buffer and Precision Plus Dual Xtra Protein Standards (Bio-Rad161-0377) were used to estimate the molecular weight of proteins. For Western blots, the SDS-PAGE separated proteins were transferred to nitrocellulose membranes. Non-specific binding to the membrane was blocked with Odyssey® PBS Blocking Buffer (Fisher Scientific NC9877369) and probed with the antibodies listed below. Blots were imaged on an Odyssey CLx and analyzed using ImageStudio2.1 (Li-COR). The following primary antibodies were used: monoclonal mouse anti-FLAG M2 (Millipore Sigma F3165), monoclonal mouse anti-strep (StrepMab) Classic iba LifeSciences 2-1507-001), custom generated Rabbit anti-Cbp1 (Rb83), custom generated Rabbit anti-RYP1, mouse anti-LAMP1 (cell signaling D401S), mouse anti-alpha-tubulin (Novus biological DM1A), Rabbit anti-calnexin (abcam 22595), mouse anti-His (ABclonal AE003).

## Abbreviations

Hc: Histoplasma capsulatum
Pb03: Paracoccidioides americana
BMDM: Bone marrow derived macrophage
ISR: Integrated Stress Response

## Acknowledgements

We thank members of the Sil, Rosenberg, and Hoving labs for helpful discussions. We would like to thank Lori Kohlstaedt, the facility director of the Vincent J. Proteomics/Mass Spectrometry Laboratory at UC Berkeley, for helpful advice and work done on this project. We thank the X-ray Crystallography Facility at University of California, San Francisco and Liam McKay for his coordination of the remote data collection. We appreciate the use of Beamline 8.3.1 at the Advanced Light Source, which is supported by the University of California Office of the President MRPI program, Plexxikon Inc. and the Integrated Diffraction Analysis Technologies program of the US Department of Energy Office of Biological and Environmental Research. The Advanced Light Source (Berkeley, CA) is a national user facility operated by Lawrence Berkeley National Laboratory on behalf of the US Department of Energy under contract number DE-AC02-05CH11231, Office of Basic Energy Sciences. We would like to thank James Holton and George Meigs for their assistance at the beamline. Finally, we thank Nelesh Govender (National Institute for Communicable Diseases, South Africa) for providing the *Emergomyces africanus* strain.

## Supporting Figure Legends

**Fig. S1. InstantBlue™ stained SDS-PAGE gel of Cbp1 homologs purified directly from culture supernatants**. Cbp1 was purified from supernatants of the indicated strains (*Hc* G217B; *Hc* G186AR; *Hc* H88; *Hc* G217B *ura5 cbp1* carrying either Pb03, *E. crescens*, or *Es. africanus* Cbp1). Purified proteins were subjected to SDS-PAGE and InstantBlue™ staining.

**Fig. S2. Comparisons of the previously published G186AR NMR structure with the H88 and Pb03 Cbp1 crystal structure**. A. Alignment of the Pb03 and H88 crystal structures with the G186AR NMR structure shows major differences in the arrangement of the helices. The RMSD scores from the alignments of G186AR to Pb03 and H88 are 5.608 and 5.879 respectively. B. Highlight of the aromatic (colored mint green) and aliphatic (colored red) residues to showcase the greasy patch formed by the C-terminal helices of each monomer and the aromatic residues packing into the hydrophobic core. C. Charge distribution over the surface of the protein structures shows major differences in the negative charge at neutral pH. Negative charge is indicated on surface with red, positive charge with blue, and neutral with white. The Pb03 dimer has a negative charge patch at the bottom of the C-terminal helical bundles, whereas the H88 dimer is only partially negative there. The H88 dimer has a negative groove that runs along the side of the protein. Only the Pb03 structure has a partial positive charge over the N-terminal helices. The electrostatic surfaces were determine using the APBS/PDB2PQR software (48, 49). D. The tertiary structures of *Emergomyces homologs* were modelled on the Pb03 Cbp1 backbone using MODELLER (50) and the surface charge distribution was determined using the APBS/PDB2PQR software.

**Fig. S3. Generation of chimeric *Hc-E. crescens* and Pb03-*E. crescens* alleles**. A. Illustration of residue changes made to *E. crescens* Cbp1 to generate chimeras with either G217B or Pb03 Cbp1. The protein alignments were compared to the alanine scan of G217B to identify residues that are required for lysis in G217B Cbp1 but not conserved in *E. crescens* Cbp1. Based on the H88 and Pb03 Cbp1 structures, we determined which of these residues were oriented outwards on the surface. These residues were also checked for whether their side chains differed in size, polarity, or charge. The proposed chimeric constructs are illustrated. The green (G217B) and yellow (Pb03) residues represent those in the N-terminal helix whereas the darker blue (G217B) and lighter blue (Pb03) residues represent those in the C-terminal loops. B. The selected differential residues that were mutated in the N-terminal helix and C-terminal loops are highlighted in green and teal respectively in the Pb03 and H88 structures.

**Fig. S4. Tunicamycin treatment results in BMDM lysis**. As a positive control for BMDM lysis, tunicamycin (Tm) was added to BMDMs at the indicated concentrations at time zero. Macrophage lysis was monitored by LDH release.

**Fig. S5. Controls for fractionation to assess rupture of small membranous compartments or *Hc*-containing phagosome**. A. BMDMs were mock-infected (uninf) or infected with either the “WT” strain (*Hc* G217B *ura5*^*-*^ carrying a *URA5* vector control), the *cbp1* mutant (*Hc* G217B *ura5*^*-*^ *cbp1*^*-*^) carrying either the vector control or G217B Cbp1 with 3XFLAG (G217B Cbp1 3XFLAG), Pb03 Cbp1 with 3XFLAG (Pb03 Cbp1 3XFLAG), WT G217B *ura5*^*-*^ carrying Yps-3 3XFLAG (Yps-3 3XFLAG), or the *Hc* G217B *ura5*^*-*^ *yps3*^*-*^ mutant (*yps3*^*-*^). Macrophage lysates were subjected to fractionation to separate cytosolic and membrane fractions. The fraction containing *Hc* yeast and macrophage nuclei with some contaminating cytosolic and membrane components was analyzed by SDS-PAGE and Western blotting using anti-Calnexin (endoplasmic reticulum), anti-α-Tubulin (cytosol), anti-FLAG or anti-Cbp1 antibodies. B. The cytosolic, membrane, and *Hc*/nuclear fractions were subjected to SDS-PAGE and Western blotting using anti-LAMP1 (marking the late endosomes and lysosomes) or anti-Ryp1 (an *Hc* transcription factor that is not released from the *Hc* cells, indicating that they are intact).

**Fig. S6. BMDM infections with *Hc* strains expressing tagged versions of *Hc* or Pb03 Cbp1**. A. BMDMs were mock-infected (uninf) or infected with the “WT” strain of *Hc* G217B *ura5*^*-*^ carrying the vector control or the *Hc* G217B *cbp1* mutant strain carrying either the *URA5* vector control or 3 independent transformants of Pb03 Cbp1 tagged with 3XFLAG. LDH release was monitored over time. B. BMDMs were mock-infected (uninf) or infected with the “WT” strain of *Hc* G217B *ura5*^*-*^ carrying the *URA5* vector control or the *Hc* G217B *cbp1* mutant strain carrying either the *URA5* vector control, untagged G217B Cbp1, or 3 independent transformants of G217B Cbp1 tagged with 3XFLAG. LDH release was monitored over time. C. BMDMs were mock-infected (uninf) or infected with the “WT” strain of *Hc* G217B *ura5*^*-*^ carrying the vector control or with the *Hc* G217B *cbp1* mutant carrying G217B Cbp1 or Pb03 Cbp1 untagged or tagged with 1XStrep or 2XStrep respectively. Other strains included *Hc* G217B *cbp1* mutant transformed with a variety of *Emergomyces* Cbp1 alleles. LDH release was monitored over time.

**Fig. S7. Determination of the optimal molar ratio for Cbp1:Yps-3 complex**. Purified Cbp1:Yps-3-6xhis were mixed in the following molar ratios: 1:1, 1:2, 2:1, and 5:1 and then Yps-3-6xhis was pulled down with Ni-NTA beads to determine if it co-purified with Cbp1.

**Fig. S8. Generation of the *yps-3* deletion mutant in *Hc* G217B using CRISPR-Cas9**. A. Schematic of CRISPR sgRNA guide target sites (triangles) for the *YPS-3* locus and the location of internal and external primers that were used to probe for the deletion of the locus in panel B. Purple box indicates the *YPS-3* coding sequence. B. PCR of *YPS-3* locus from genomic DNA from WT *Hc* G217B and *yps-3*Δ mutant utilizing the external or internal primers designated in A.

**S9 Table:**
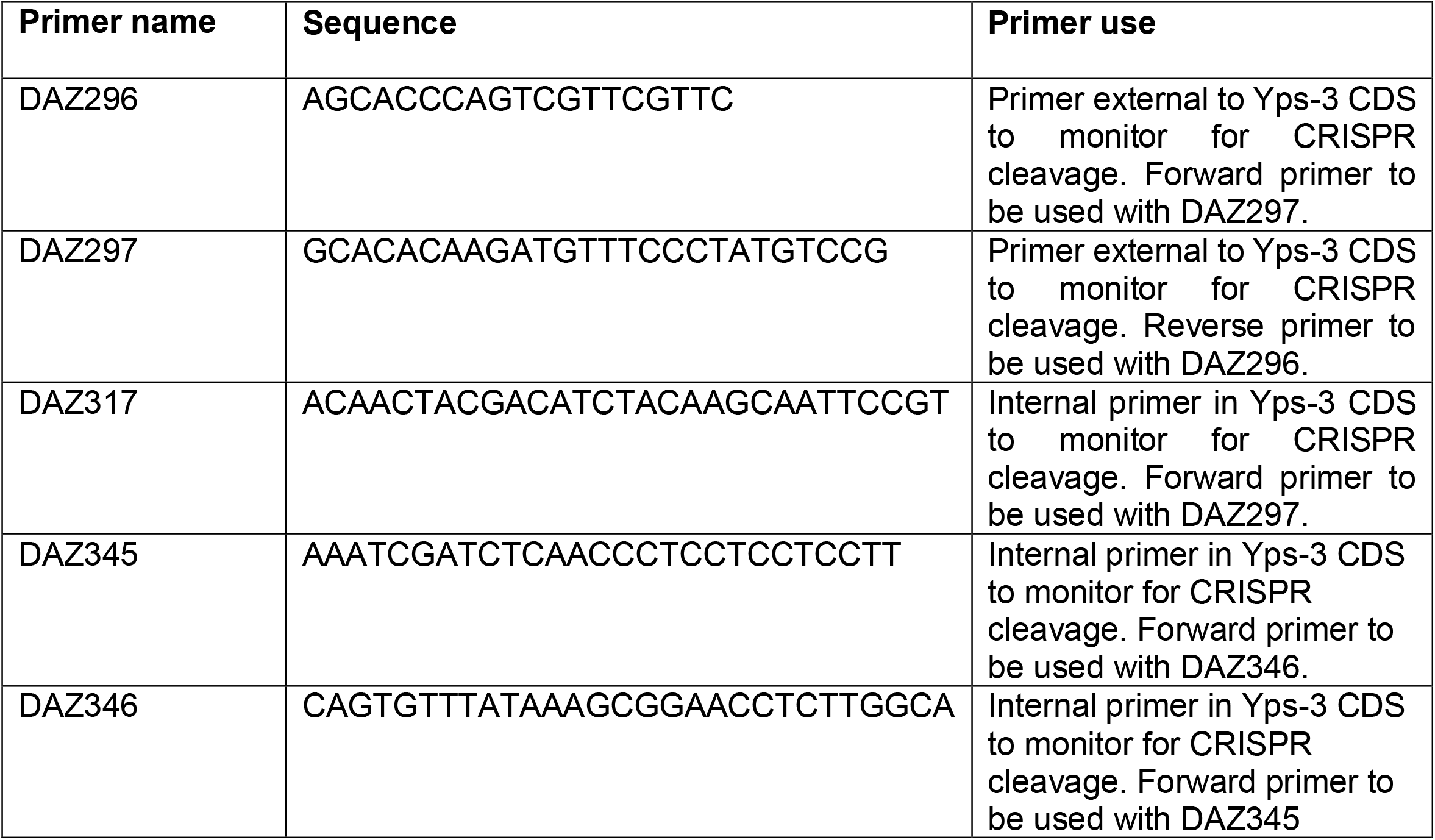
Primers used in this study.

## References

1. Alto NM, Orth K. Subversion of cell signaling by pathogens. Cold Spring Harb Perspect Biol. 2012;4(9):a006114.

2. Ham H, Sreelatha A, Orth K. Manipulation of host membranes by bacterial effectors. Nat Rev Microbiol. 2011;9(9):635–46.

3. Hakimi MA, Bougdour A. Toxoplasma’s ways of manipulating the host transcriptome via secreted effectors. Curr Opin Microbiol. 2015;26:24–31.

4. Mitchell G, Chen C, Portnoy DA. Strategies Used by Bacteria to Grow in Macrophages. Microbiol Spectr. 2016;4(3).

5. Pradhan A, Ghosh S, Sahoo D, Jha G. Fungal effectors, the double edge sword of phytopathogens. Curr Genet. 2021;67(1):27–40.

6. Asrat S, de Jesus DA, Hempstead AD, Ramabhadran V, Isberg RR. Bacterial pathogen manipulation of host membrane trafficking. Annu Rev Cell Dev Biol. 2014;30:79–109.

7. Fredlund J, Enninga J. Cytoplasmic access by intracellular bacterial pathogens. Trends Microbiol. 2014;22(3):128–37.

8. Personnic N, Barlocher K, Finsel I, Hilbi H. Subversion of Retrograde Trafficking by Translocated Pathogen Effectors. Trends Microbiol. 2016;24(6):450–62.

9. Brown GD, Denning DW, Levitz SM. Tackling human fungal infections. Science. 2012;336(6082):647.

10. Brown GD, Denning DW, Gow NA, Levitz SM, Netea MG, White TC. Hidden killers: human fungal infections. Sci Transl Med. 2012;4(165):165rv13.

11. Chu JH, Feudtner C, Heydon K, Walsh TJ, Zaoutis TE. Hospitalizations for endemic mycoses: a population-based national study. Clin Infect Dis. 2006;42(6):822–5.

12. Arauz AB, Papineni P. Histoplasmosis. Infect Dis Clin North Am. 2021;35(2):471–91.

13. Kauffman CA. Histoplasmosis: a clinical and laboratory update. Clin Microbiol Rev. 2007;20(1):115–32.

14. Woods JP. Revisiting old friends: Developments in understanding Histoplasma capsulatum pathogenesis. J Microbiol. 2016;54(3):265–76.

15. Lockhart SR, Toda M, Benedict K, Caceres DH, Litvintseva AP. Endemic and Other Dimorphic Mycoses in The Americas. J Fungi (Basel). 2021;7(2).

16. Benedict K, Mody RK. Epidemiology of Histoplasmosis Outbreaks, United States, 1938-2013. Emerg Infect Dis. 2016;22(3):370–8.

17. Maiga AW, Deppen S, Scaffidi BK, Baddley J, Aldrich MC, Dittus RS, et al. Mapping Histoplasma capsulatum Exposure, United States. Emerg Infect Dis. 2018;24(10):1835–9.

18. Garfoot AL, Rappleye CA. Histoplasma capsulatum surmounts obstacles to intracellular pathogenesis. FEBS J. 2016;283(4):619–33.

19. Holbrook ED, Rappleye CA. Histoplasma capsulatum pathogenesis: making a lifestyle switch. Curr Opin Microbiol. 2008;11(4):318–24.

20. Maresca B, Kobayashi GS. Dimorphism in Histoplasma capsulatum: a model for the study of cell differentiation in pathogenic fungi. Microbiol Rev. 1989;53(2):186–209.

21. Howard DH. Intracellular Behavior of Histoplasma Capsulatum. J Bacteriol. 1964;87:33–8.

22. Howard DH. Intracellular Growth of Histoplasma Capsulatum. J Bacteriol. 1965;89:518–23.

23. Newman SL, Bucher C, Rhodes J, Bullock WE. Phagocytosis of Histoplasma capsulatum yeasts and microconidia by human cultured macrophages and alveolar macrophages. Cellular cytoskeleton requirement for attachment and ingestion. J Clin Invest. 1990;85(1):223–30.

24. Shen Q, Rappleye CA. Differentiation of the fungus Histoplasma capsulatum into a pathogen of phagocytes. Curr Opin Microbiol. 2017;40:1–7.

25. Newman SL, Gootee L, Hilty J, Morris RE. Human macrophages do not require phagosome acidification to mediate fungistatic/fungicidal activity against Histoplasma capsulatum. J Immunol. 2006;176(3):1806–13.

26. Strasser JE, Newman SL, Ciraolo GM, Morris RE, Howell ML, Dean GE. Regulation of the macrophage vacuolar ATPase and phagosome-lysosome fusion by Histoplasma capsulatum. J Immunol. 1999;162(10):6148–54.

27. Batanghari JW, Goldman WE. Calcium dependence and binding in cultures of Histoplasma capsulatum. Infect Immun. 1997;65(12):5257–61.

28. Batanghari JW, Deepe GS, Jr., Di Cera E, Goldman WE. Histoplasma acquisition of calcium and expression of CBP1 during intracellular parasitism. Mol Microbiol. 1998;27(3):531–9.

29. Sebghati TS, Engle JT, Goldman WE. Intracellular parasitism by Histoplasma capsulatum: fungal virulence and calcium dependence. Science. 2000;290(5495):1368–72.

30. Kugler S, Young B, Miller VL, Goldman WE. Monitoring phase-specific gene expression in Histoplasma capsulatum with telomeric GFP fusion plasmids. Cell Microbiol. 2000;2(6):537–47.

31. Beck MR, Dekoster GT, Cistola DP, Goldman WE. NMR structure of a fungal virulence factor reveals structural homology with mammalian saposin B. Mol Microbiol. 2009;72(2):344–53.

32. Beck MR, DeKoster GT, Hambly DM, Gross ML, Cistola DP, Goldman WE. Structural features responsible for the biological stability of Histoplasma’s virulence factor CBP. Biochemistry. 2008;47(15):4427–38.

33. Isaac DT, Berkes CA, English BC, Murray DH, Lee YN, Coady A, et al. Macrophage cell death and transcriptional response are actively triggered by the fungal virulence factor Cbp1 during H. capsulatum infection. Mol Microbiol. 2015;98(5):910–29.

34. English BC, Van Prooyen N, Ord T, Ord T, Sil A. The transcription factor CHOP, an effector of the integrated stress response, is required for host sensitivity to the fungal intracellular pathogen Histoplasma capsulatum. PLoS Pathog. 2017;13(9):e1006589.

35. Bohse ML, Woods JP. RNA interference-mediated silencing of the YPS3 gene of Histoplasma capsulatum reveals virulence defects. Infect Immun. 2007;75(6):2811–7.

36. Bohse ML, Woods JP. Surface localization of the Yps3p protein of Histoplasma capsulatum. Eukaryot Cell. 2005;4(4):685–93.

37. Bohse ML, Woods JP. Expression and interstrain variability of the YPS3 gene of Histoplasma capsulatum. Eukaryot Cell. 2007;6(4):609–15.

38. Schwartz IS, Govender NP, Sigler L, Jiang Y, Maphanga TG, Toplis B, et al. Emergomyces: The global rise of new dimorphic fungal pathogens. PLoS Pathog. 2019;15(9):e1007977.

39. Govender NP, Grayson W. Emergomycosis (Emergomyces africanus) in Advanced HIV Disease. Dermatopathology (Basel). 2019;6(2):63–9.

40. Samaddar A, Sharma A. Emergomycosis, an Emerging Systemic Mycosis in Immunocompromised Patients: Current Trends and Future Prospects. Front Med (Lausanne). 2021;8:670731.

41. Dukik K, Munoz JF, Jiang Y, Feng P, Sigler L, Stielow JB, et al. Novel taxa of thermally dimorphic systemic pathogens in the Ajellomycetaceae (Onygenales). Mycoses. 2017;60(5):296–309.

42. McBride JA, Gauthier GM, Klein BS. Turning on virulence: Mechanisms that underpin the morphologic transition and pathogenicity of Blastomyces. Virulence. 2019;10(1):801–9.

43. Sil A, Andrianopoulos A. Thermally Dimorphic Human Fungal Pathogens--Polyphyletic Pathogens with a Convergent Pathogenicity Trait. Cold Spring Harb Perspect Med. 2014;5(8):a019794.

44. Yang Z. PAML 4: phylogenetic analysis by maximum likelihood. Mol Biol Evol. 2007;24(8):1586–91.

45. Holbrook ED, Edwards JA, Youseff BH, Rappleye CA. Definition of the extracellular proteome of pathogenic-phase Histoplasma capsulatum. J Proteome Res. 2011;10(4):1929–43.

46. Holm L. DALI and the persistence of protein shape. Protein Sci. 2020;29(1):128–40.

47. Madej T, Lanczycki CJ, Zhang D, Thiessen PA, Geer RC, Marchler-Bauer A, et al. MMDB and VAST+: tracking structural similarities between macromolecular complexes. Nucleic Acids Res. 2014;42(Database issue):D297–303.

48. Baker NA, Sept D, Joseph S, Holst MJ, McCammon JA. Electrostatics of nanosystems: application to microtubules and the ribosome. Proc Natl Acad Sci U S A. 2001;98(18):10037–41.

49. Dolinsky TJ, Nielsen JE, McCammon JA, Baker NA. PDB2PQR: an automated pipeline for the setup of Poisson-Boltzmann electrostatics calculations. Nucleic Acids Res. 2004;32(Web Server issue):W665–7.

50. Sali A, Blundell TL. Comparative protein modelling by satisfaction of spatial restraints. J Mol Biol. 1993;234(3):779–815.

51. Taylor ML, Espinosa-Schoelly ME, Iturbe R, Rico B, Casasola J, Goodsaid F. Evaluation of phagolysosome fusion in acridine orange stained macrophages infected with Histoplasma capsulatum. Clin Exp Immunol. 1989;75(3):466–70.

52. Eissenberg LG, Goldman WE, Schlesinger PH. Histoplasma capsulatum modulates the acidification of phagolysosomes. J Exp Med. 1993;177(6):1605–11.

53. Kujoth GC, Sullivan TD, Klein BS. Gene Editing in Dimorphic Fungi Using CRISPR/Cas9. Curr Protoc Microbiol. 2020;59(1):e132.

54. Joehnk B, Voorhies, M., Walcott, K., Sil, A. Recyclable CRISPR/Cas9 mediated gene disruption and deletions in Histoplasma, manuscript in preparation.

55. Baltazar LM, Zamith-Miranda D, Burnet MC, Choi H, Nimrichter L, Nakayasu ES, et al. Concentration-dependent protein loading of extracellular vesicles released by Histoplasma capsulatum after antibody treatment and its modulatory action upon macrophages. Sci Rep. 2018;8(1):8065.

56. Alves LR, Peres da Silva R, Sanchez DA, Zamith-Miranda D, Rodrigues ML, Goldenberg S, et al. Extracellular Vesicle-Mediated RNA Release in Histoplasma capsulatum. mSphere. 2019;4(2).

57. Finkel-Jimenez B, Wuthrich M, Klein BS. BAD1, an essential virulence factor of Blastomyces dermatitidis, suppresses host TNF-alpha production through TGF-beta-dependent and -independent mechanisms. J Immunol. 2002;168(11):5746–55.

58. Brandhorst T, Wuthrich M, Finkel-Jimenez B, Klein B. A C-terminal EGF-like domain governs BAD1 localization to the yeast surface and fungal adherence to phagocytes, but is dispensable in immune modulation and pathogenicity of Blastomyces dermatitidis. Mol Microbiol. 2003;48(1):53–65.

59. Rooney PJ, Klein BS. Sequence elements necessary for transcriptional activation of BAD1 in the yeast phase of Blastomyces dermatitidis. Eukaryot Cell. 2004;3(3):785–94.

60. Beaussart A, Brandhorst T, Dufrene YF, Klein BS. Blastomyces Virulence Adhesin-1 Protein Binding to Glycosaminoglycans Is Enhanced by Protein Disulfide Isomerase. mBio. 2015;6(5):e01403–15.

61. da Silva TA, Roque-Barreira MC, Casadevall A, Almeida F. Extracellular vesicles from Paracoccidioides brasiliensis induced M1 polarization in vitro. Sci Rep. 2016;6:35867.

62. Schwartz IS, Kenyon C, Lehloenya R, Claasens S, Spengane Z, Prozesky H, et al. AIDS-Related Endemic Mycoses in Western Cape, South Africa, and Clinical Mimics: A Cross-Sectional Study of Adults With Advanced HIV and Recent-Onset, Widespread Skin Lesions. Open Forum Infect Dis. 2017;4(4):ofx186.

63. Eddy SR. Accelerated Profile HMM Searches. PLoS Comput Biol. 2011;7(10):e1002195.

64. Altschul SF, Madden TL, Schaffer AA, Zhang J, Zhang Z, Miller W, et al. Gapped BLAST and PSI-BLAST: a new generation of protein database search programs. Nucleic Acids Res. 1997;25(17):3389–402.

65. Do CB, Mahabhashyam MS, Brudno M, Batzoglou S. ProbCons: Probabilistic consistency-based multiple sequence alignment. Genome Res. 2005;15(2):330–40.

66. Nguyen LT, Schmidt HA, von Haeseler A, Minh BQ. IQ-TREE: a fast and effective stochastic algorithm for estimating maximum-likelihood phylogenies. Mol Biol Evol. 2015;32(1):268–74.

67. Staff PNTD. Correction: Molecular detection of airborne Emergomyces africanus, a thermally dimorphic fungal pathogen, in Cape Town, South Africa. PLoS Negl Trop Dis. 2018;12(5):e0006468.

68. Schwartz IS, McLoud JD, Berman D, Botha A, Lerm B, Colebunders R, et al. Molecular detection of airborne Emergomyces africanus, a thermally dimorphic fungal pathogen, in Cape Town, South Africa. PLoS Negl Trop Dis. 2018;12(1):e0006174.

