## Supplementary figures and images for "Cbp1, a rapidly evolving fungal virulence factor, forms an effector complex that drives macrophage lysis"

### Supp_Figure_1

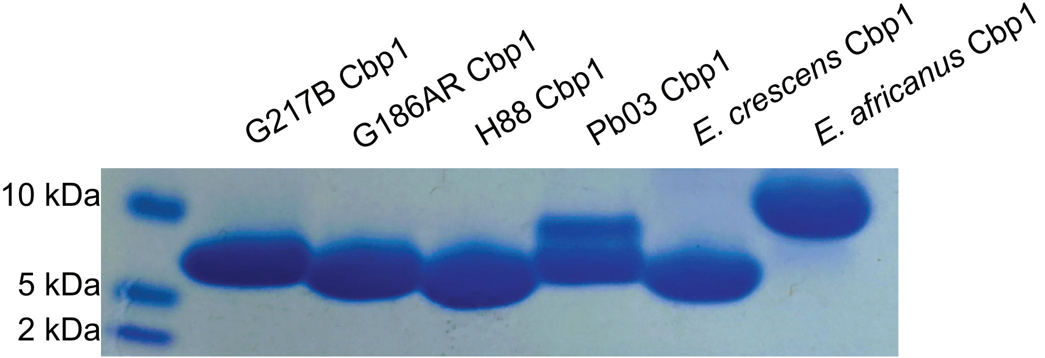

### Supp_Figure_2

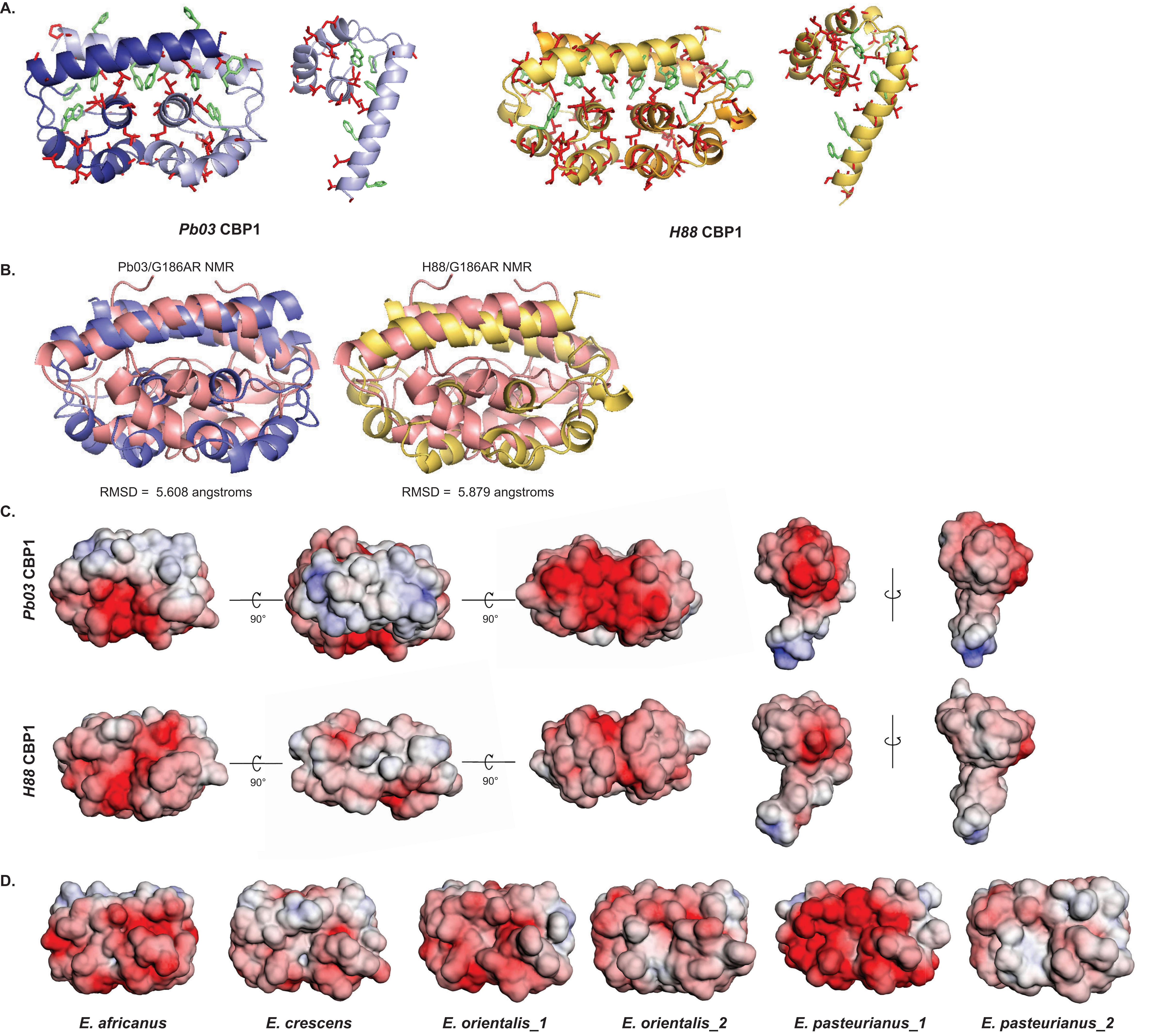

### Supp_Figure_3

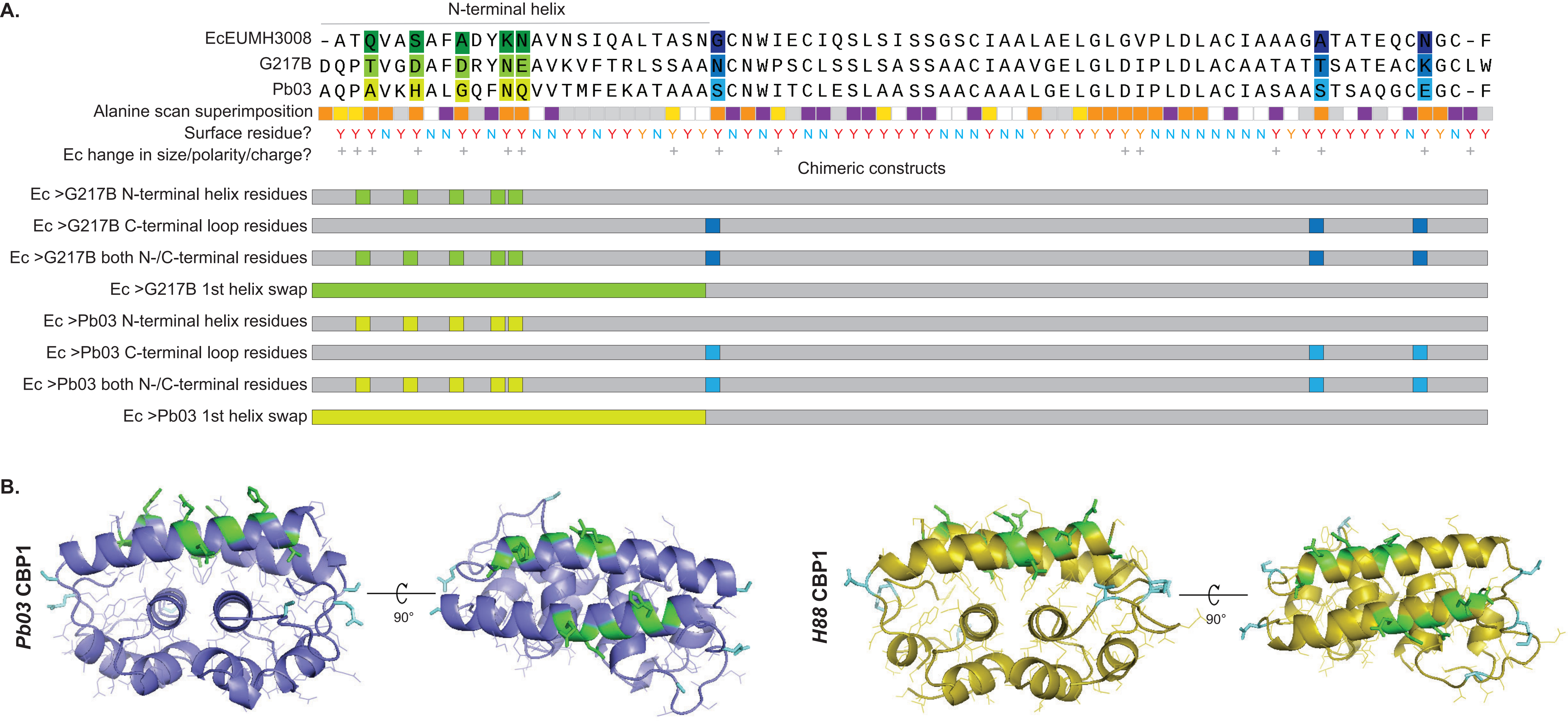

### Supp_Figure_4

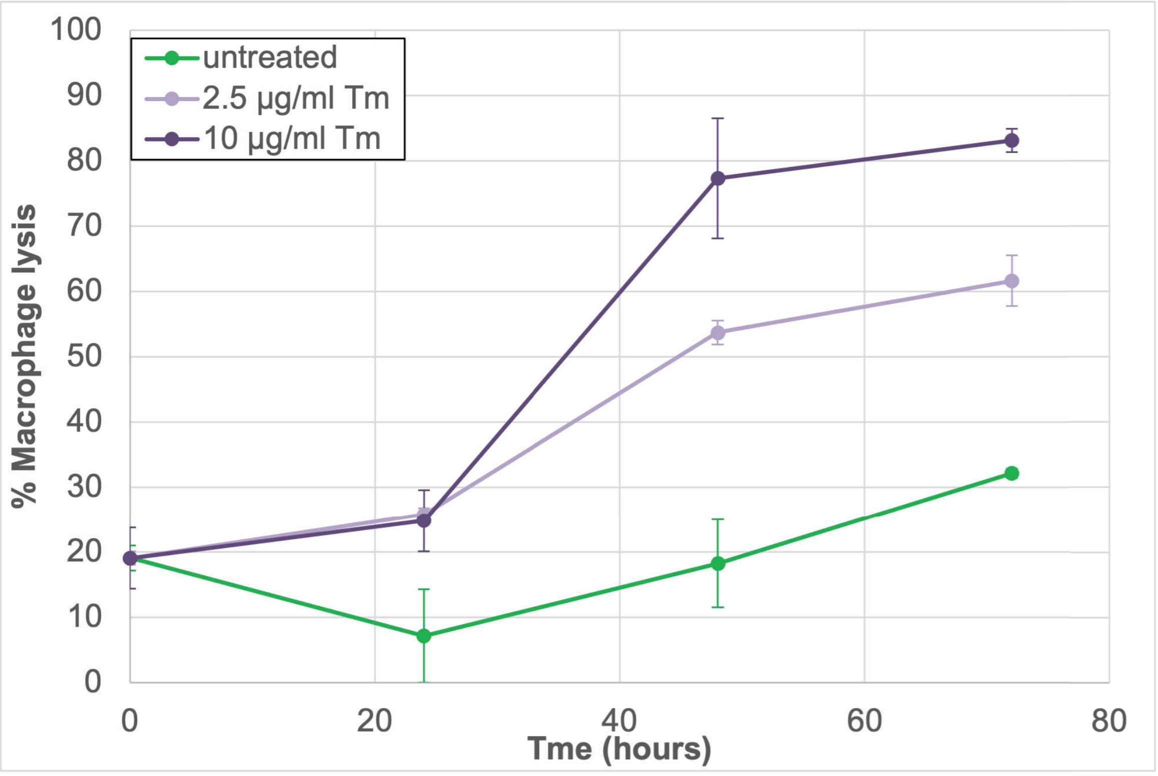

### Supp_Figure_5

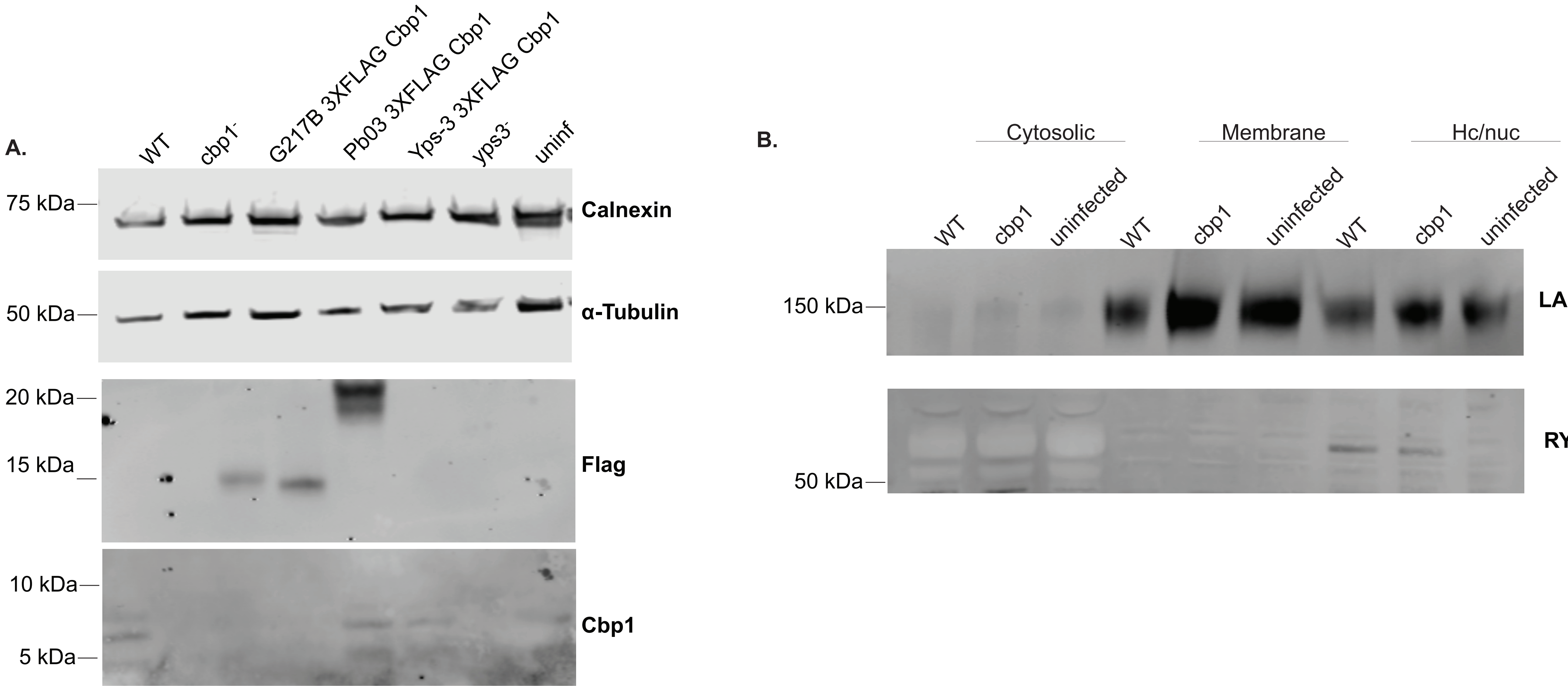

### Supp_Figure_6

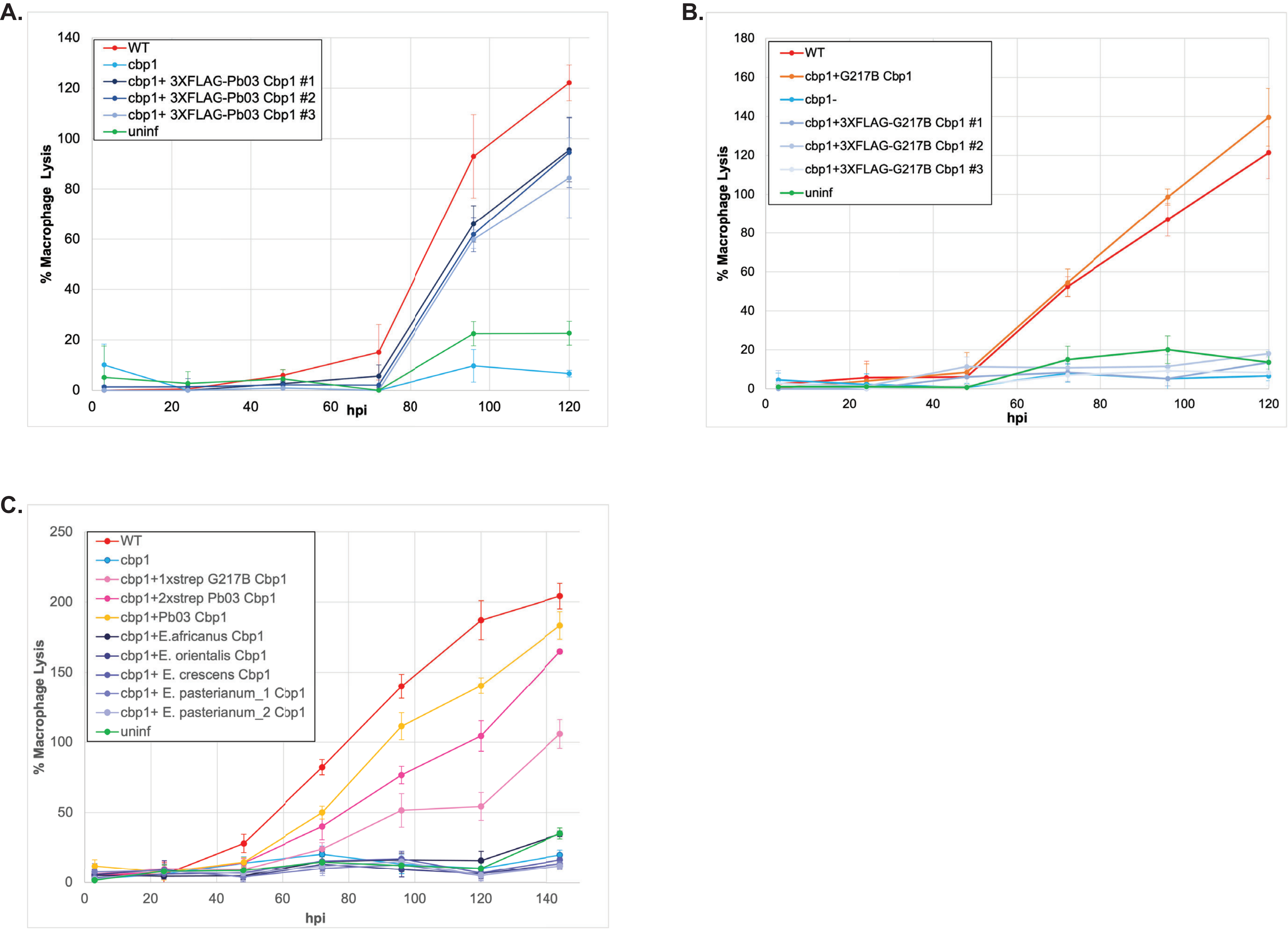

### Supp_Figure_7

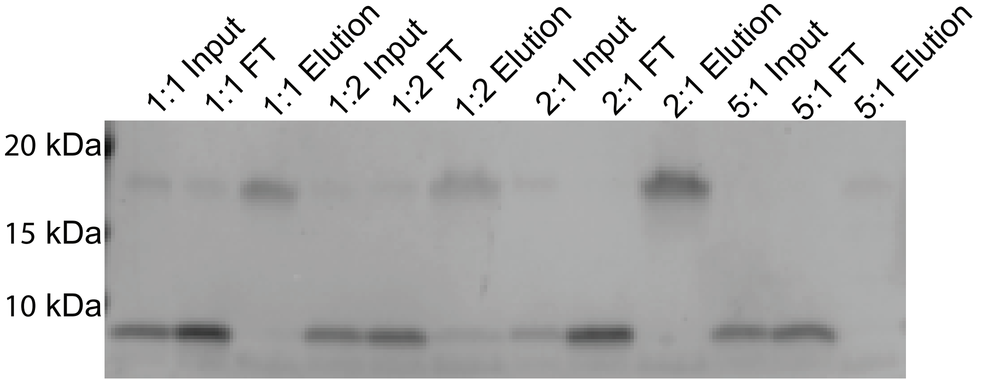

### Supp_Figure_8

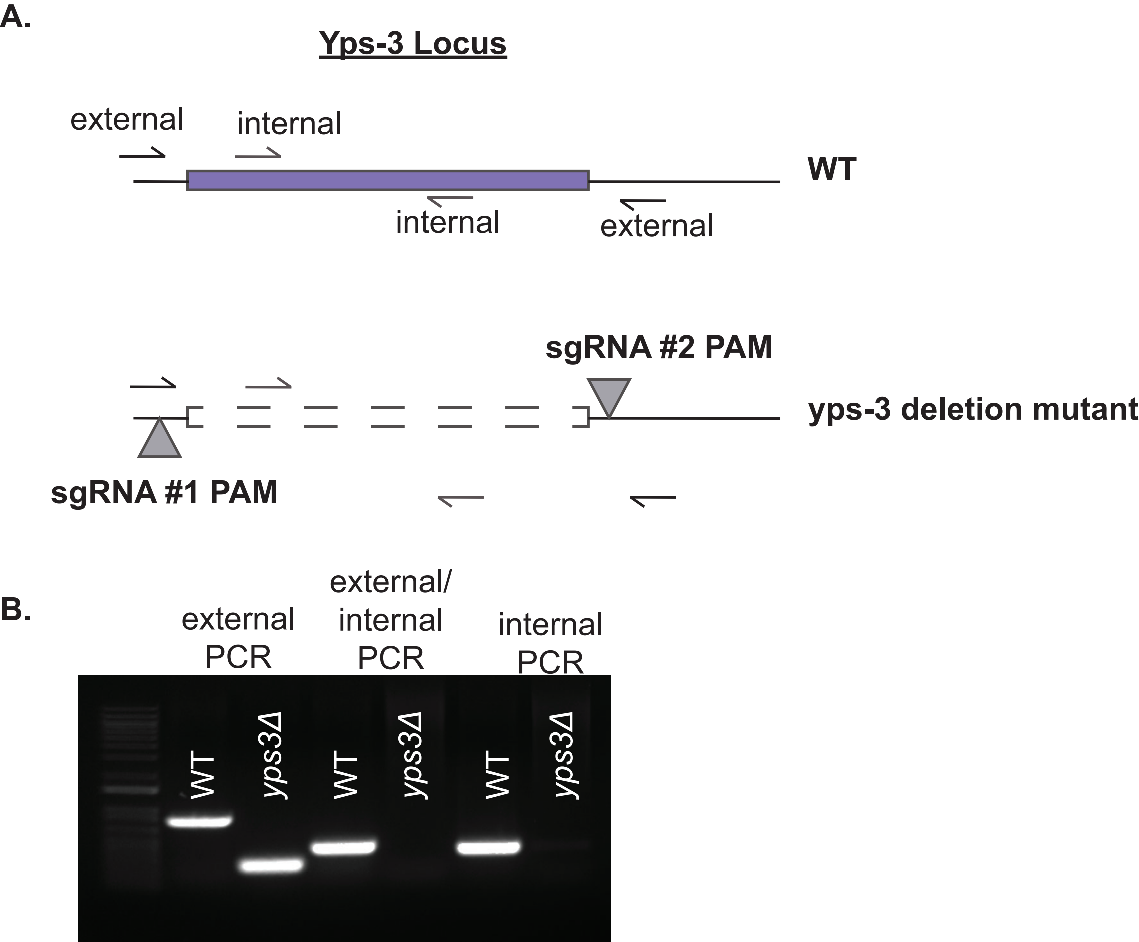
